# TRIB2 couples UCP1 degradation to thermogenic adaptation and metabolic health

**DOI:** 10.1101/2025.10.26.684708

**Authors:** Jiin Horng Lee, Li-Yun Chueh, Sui-Chih Tien, Hsiao-lin Lee, Kou-Ray Lin, Tung-Yuan Lee, Tony Pan-Hou Che, Jeffrey J.Y. Yen, Chun-Mei Hu, Yi-Cheng Chang

## Abstract

Genetic variation in TRIB2 has been associated with thermogenesis and fat accumulation, yet the underlying molecular mechanisms remain unclear. Uncoupling protein 1 (UCP1) is a key regulator of adaptive thermogenesis and energy expenditure. Here, we identified a human TRIB2 variant associated with migration patterns along latitude and environmental temperature, which destabilized its mRNA and is consistent with enhanced thermogenic capacity. We demonstrate that loss of *Trib2* protects mice from diet-induced obesity, alleviates hepatic steatosis, and improves glucose tolerance and insulin sensitivity. *Trib2* knockout mice exhibit elevated UCP1 expression in brown adipose tissue, leading to enhanced thermogenesis and increased heat production. Mechanistically, TRIB2 acts as a scaffold that binds UCP1 through its pseudokinase domain and recruits the E3 ligase MYCBP2, thereby promoting UCP1 ubiquitination and proteasomal degradation. Collectively, our findings reveal a previously unrecognized post-translational mechanism that regulates UCP1 stability and link TRIB2 function to both thermogenic adaptation and metabolic health, thereby highlighting TRIB2 as a potential therapeutic target for obesity and metabolic disorders.

## Introduction

Obesity, characterized by excess body fat(World Health Organization, 2023) with detrimental effects on health, is typically defined by a body mass index (BMI) of 30 or higher(World Health Organization). This condition arises from an imbalance between calorie intake and expenditure(Hill & Commerford, 1996), influenced by genetics(Pulit *et al*, 2019; Yengo *et al*, 2018), environment(Nicolaidis, 2019), metabolism(Parati *et al*, 2007; Singla *et al*, 2010), and behavior(Bray, 2004). Metabolic syndrome often accompanies obesity and is associated with various health problems, including chronic diseases such as type 2 diabetes(Alexopoulos *et al*, 2014; Klein *et al*, 2022), cardiovascular diseases(Canoy, 2008) (including ischemic heart disease, heart failure, and stroke), hypertension(Hall *et al*, 2015), and certain cancers(Harvey *et al*, 2023; Lee *et al*, 2019; Mandic *et al*, 2023)(such as breast, colon, and endometrial cancer). The continued rise in prevalence, along with its related health and economic impacts, significantly strains the healthcare system(Tremmel *et al*, 2017).

Brown adipose tissue (BAT) is rich in mitochondria (Enerbäck, 2009) and distinguished by the expression of uncoupling protein 1 (UCP1)(Ricquier & Bouillaud, 2000). BAT plays a crucial role in non-shivering thermogenesis (Himms-Hagen, 1984), the process by which the body produces heat and energy metabolism in the body(CANNON & NEDERGAARD, 2004; Saito *et al*, 2009). In recent years, positron emission tomography (PET) scanning has allowed for non-invasive imaging of BAT located in the neck, upper back, and between the shoulder blades of the adult human body, not only in infants and young children(Cypess *et al*, 2009; Lichtenbelt *et al*, 2009; Saito *et al*., 2009; Seki *et al*, 2022; Yoneshiro *et al*, 2011). BAT activity can be stimulated by various factors, including cold exposure(Lichtenbelt *et al*., 2009; Orava *et al*, 2011), certain hormones(Boström *et al*, 2012; Orava *et al*., 2011), and drugs(Cypess *et al*, 2015). The increasing BAT activity may have beneficial effects on metabolism and body weight regulation (Cypess *et al*., 2009; Hachemi & M, 2023; Harb *et al*, 2023; Lichtenbelt *et al*., 2009; Scheele & Nielsen, 2017), leading to increased energy expenditure, improved glucose metabolism, and reduced adiposity. There is considerable interest in developing therapies that target BAT to attenuate obesity and metabolic disorders.

A multi-ethnic genome-wide meta-analysis of ectopic fat accumulation identified a single-nucleotide variant (SNV), rs10198628, in the TRIB2 gene, which is associated with pericardial fat mas(Chu *et al*, 2017). This finding was validated in an independent cohort of 5,487 individuals of European ancestry(Fox *et al*, 2012). A genome-wide association study (GWAS) confirmed the TRIB2 variant, rs1057001 in the 3’ untranslated region (UTR), associated with waist circumference in 2,958 Chinese(Wang *et al*, 2016). This variant, associated with BMI and waist circumference, was confirmed in another study involving 3,013 Japanese individuals, with a high prevalence of the derived A allele in East Asians(Nakayama *et al*, 2013). Interestingly, the rs1057001 A allele of this variant was associated with higher expression levels of thermogenic genes in human subcutaneous and visceral adipose tissues(Nakayama & Iwamoto, 2017). These findings suggest that environmental or physiological factors, particularly temperature-related thermogenic demands, may have exerted positive selective pressure on TRIB2 during human adaptation in East Asians.

Tribbles homolog 2 (TRIB2) is a member of the CAMK/serine/threonine pseudokinase family(Foulkes *et al*, 2015; Jamieson *et al*, 2022; Shrestha *et al*, 2020). Structurally, TRIB2 consists of three main domains: an N-terminal PEST domain, a central pseudokinase domain, and a C-terminal MAPK (mitogen-activated protein kinase) E3 ligase-binding domain(Qiao *et al*, 2013; Rechsteiner & Rogers, 1996; Rogers *et al*, 1986; Wang *et al*, 2013). Although the central pseudokinase domain of TRIB2 exhibits low ATP affinity and kinase activity (Bailey *et al*, 2015; Murphy *et al*, 2015; Murphy *et al*, 2014), it still maintains the ability to interact with several signaling proteins and function as a scaffold protein(Dobens Jr. & Bouyain, 2012). TRIB2 mediates multiple signaling pathways in various cancers, such as promoting degradation of CCAAT/enhancer-binding protein alpha (C/EBP-α) in acute myeloid leukemia (AML)(Dedhia *et al*, 2010; Keeshan *et al*, 2006), repressing Forkhead box O (FOXO) in melanoma(Zanella *et al*, 2010), and participating in E3 ligase recruitment and Wnt/β-catenin/TCF4 signaling in response to oxidative stress in liver and lung cancers(Grandinetti *et al*, 2011; Guo *et al*, 2021; Wang *et al*., 2013; Xiang *et al*, 2021; Xu *et al*, 2014). In addition, TRIB2 inhibits 3T3L1 white pre-adipocyte differentiation by suppressing AKT activation and C/EBP-β and δ(Naiki *et al*, 2007), indicating its possible role in adipocyte differentiation. A recent study shows TRIB2 modulates AMPK activity to promote hepatic insulin resistance(Wang *et al*, 2024). Taken together, TRIB2 modulates fat distribution and metabolic health, encompassing ectopic and pericardial fat accumulation, waist circumference, and BMI by regulating adipocyte differentiation and hepatic insulin resistance. Apart from these established pathways, we hypothesize that TRIB2 may contribute to fat distribution and metabolic health by modulating thermogenic gene expression.

In this study, we demonstrate that *Trib2* knockout prevents weight gain and fat accumulation, reduces hepatic steatosis, and improves glucose tolerance and insulin sensitivity in mice. *Trib2* knockout (KO) mice exhibit elevated UCP1 expression in brown adipose tissue, resulting in enhanced energy expenditure and thermogenesis. Mechanistically, TRIB2 functions as a scaffold protein that binds UCP1 via its pseudokinase domain and recruits the MYCBP2 E3 ligase, promoting UCP1 ubiquitination and subsequent degradation.

## Results

### The *Trib2* rs1057001 A allele is associated with thermogenic adaptation and reduces TRIB2 mRNA stability

Several studies have shown that SNPs in TRIB2 are associated with subcutaneous and visceral adipose tissue(Chu *et al*., 2017; Fox *et al*., 2012; Nakayama & Iwamoto, 2017; Nakayama *et al*., 2013; Wang *et al*., 2016). The A allele of SNP rs1057001, located in the 3’UTR of the TRIB2 gene, is associated with elevated expression of thermogenic genes in adipose tissue in humans and reduced visceral fat mass(Nakayama & Iwamoto, 2017). Another SNP, rs16350, a dinucleotide deletion in absolute linkage disequilibrium (LD) with rs1057001, was located 7 bp upstream of the polyadenylation site of the human TRIB2 gene(Nakayama *et al*., 2013). These 2 SNPs, rs1057001 T > A and rs16350 ATC > A alleles, constitute a haplotype(Nakayama & Iwamoto, 2017; Nakayama *et al*., 2013). We collected a total of 17 population or subpopulation of SNP rs1057001 data from the 1000 Genomes(Auton *et al*, 2015), gnomAD v4-Genomes(Karczewski *et al*, 2020), Human Genome Diversity Project(Bergström *et al*, 2020), Siberian project(Wong *et al*, 2017), and 38KJPN(Tadaka *et al*, 2019) (Fig. 1A). We used allele frequency pie charts on a climate geographic map at the location of each population (Fig. 1B). A Similar result was observed with Nakayama, Kazuhiro et al.(Nakayama *et al*., 2013), a gradual change in the allele frequency of rs1057001 along a geographic axis suggests historical migration or gene flow associated with latitude and temperature, with the A allele being rare in sub-Saharan African regions but prevalent in East Asian and Siberian, indicative of positive natural selection during human migration. We further used Hudson’s F_ST_ to estimate the differentiation of SNP rs1057001 in 16 populations. The F_ST_ score matrices of the 16 populations were all > 0.25, indicating a very high degree of differentiation and strong genetic separation among the populations, which may be caused by specific environmental selection. (Fig. 1C). We hypothesized that the rs1057001 T > A and rs16350 ATC > A variants may affect RNA stability of TRIB2. To test this, we cloned the 3’UTR of TRIB2 from human genomic DNA into the pmirGLO dual-luciferase reporter vector. Two constructs representing different haplotypes were generated and confirmed by Sanger sequencing (Fig. 1D). The rs1057001T > A and rs16350 ATC > A haplotype is associated with decreased luciferin expression (Fig. 1E), suggesting this haplotype (Hap2) causes RNA instability, subsequently downregulating protein expression. Hence, SNP rs1057001, which was highly differentiated across different populations and caused TRIB2 RNA instability, was associated with human latitude or temperature.

**Figure 1.**
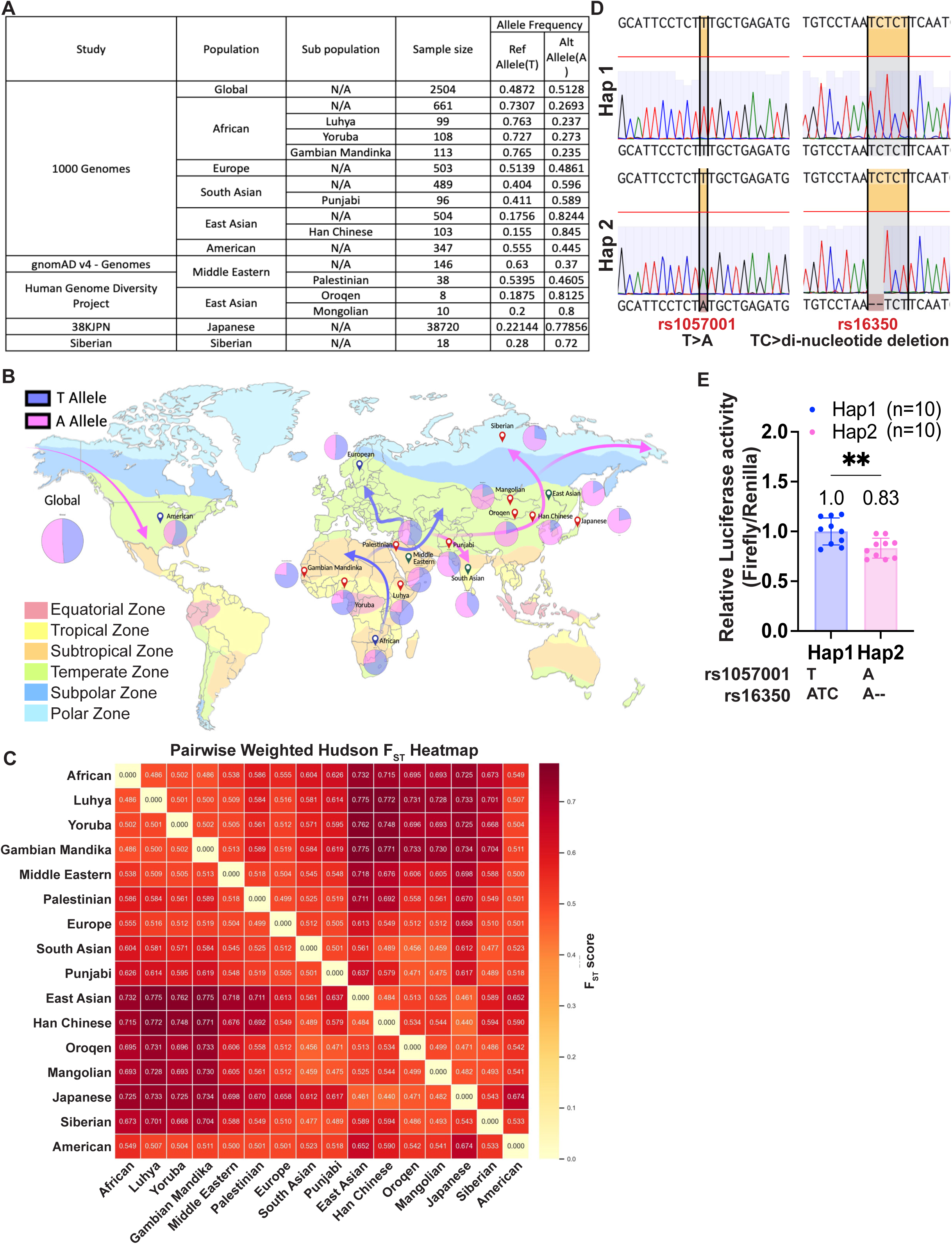
The Trib2 rs1057001 A allele is associated with thermogenic adaptation and reduces TRIB2 mRNA stability. (A) A total of 17 populations of SNP rs1057001 data from the 5 studies. (B) Allele frequency pie charts on a climate geographic map at the location of each population suggest historical migration or gene flow along that path. (C) The Hudson F_ST_ score matrix in 16 populations were >0.25, suggesting very high differentiation. (D) Hap1 rs1057001 is T, and rs16350 is TC; Hap2 rs1057001 is a T-to-A substitution, and rs16350 shows a deletion of TC. (E) The SNPs rs1057001 and rs15350 linkage disequilibrium block decreases the luciferin expression. Data B was analyzed by student unpaired t-tests. All data represent the mean ± SD. \**p* < 0.05, \*\**p* < 0.01; \*\*\**p* < 0.001; \*\*\*\**p* < 0.0001.

### Loss of *Trib2* prevents diet-induced obesity and improves glucose homeostasis in mice

We hypothesize that the loss of TRIB2 may contribute to altered fat distribution and impaired metabolic health by modulating thermogenic gene expression. To investigate this mechanism, we generated *Trib2* knockout mice (Fig EV 1A&B). Both *Trib2* wild-type (WT) and knockout (KO) were fed either the chow diet (CD) or high-fat-high-sucrose diet (HFHSD). The body weight of KO mice was significantly less than that of WT mice, both fed with CD and HFHSD (Fig. 2A). Head circumference and femur length did not differ among the four groups, suggesting that the knockout of *Trib2* does not affect mouse growth (Fig EV 1C). We measured the body composition of these four groups of mice. Under CD, there is no difference in body composition between WT and KO mice. Under HFHSD, WT mice had increased body fat and reduced lean mass compared to the KO mice. (Fig. 2B). On HFHSD, *Trib2* KO mice had reduced adipose tissue and liver weight compared to the WT HFHSD mice (Fig. 2C&D). H&E staining of perigonadal adipose tissue reveals that *Trib2* KO mice had smaller and more numerous adipocytes compared to the WT under HFHSD feeding (Fig. 2E&F). Besides that, *Trib2* KO mice had less hepatocellular steatosis compared to the WT mice on HFHSD (Fig EV 2A). The level of hepatic triglyceride (TG) content was significantly decreased in KO mice compared to the WT (Fig EV 2B). We further conducted the intraperitoneal glucose tolerance test and intraperitoneal insulin tolerance test. On HFHSD, *Trib2* KO mice displayed significantly decreased glucose tolerance and insulin resistance compared to the WT mice (Fig EV 2C&D, left panel). *Trib2* KO mice fed with HFHSD had lower serum levels of triglyceride and lower adiponectin than WT control (Fig EV 2E). We observed crown-like structures in *Trib2* WT mice, but not in KO mice, when fed a HFHSD, as revealed by F4/80 immunohistochemical staining, indicating that *Trib2* knockout protects against obesity-induced macrophage infiltration (Fig EV 2F). Thus, knockout of *Trib2* in mice protected against diet-induced weight gain and fat accumulation, attenuated hepatic steatosis, and improved glucose homeostasis.

**Figure 2.**
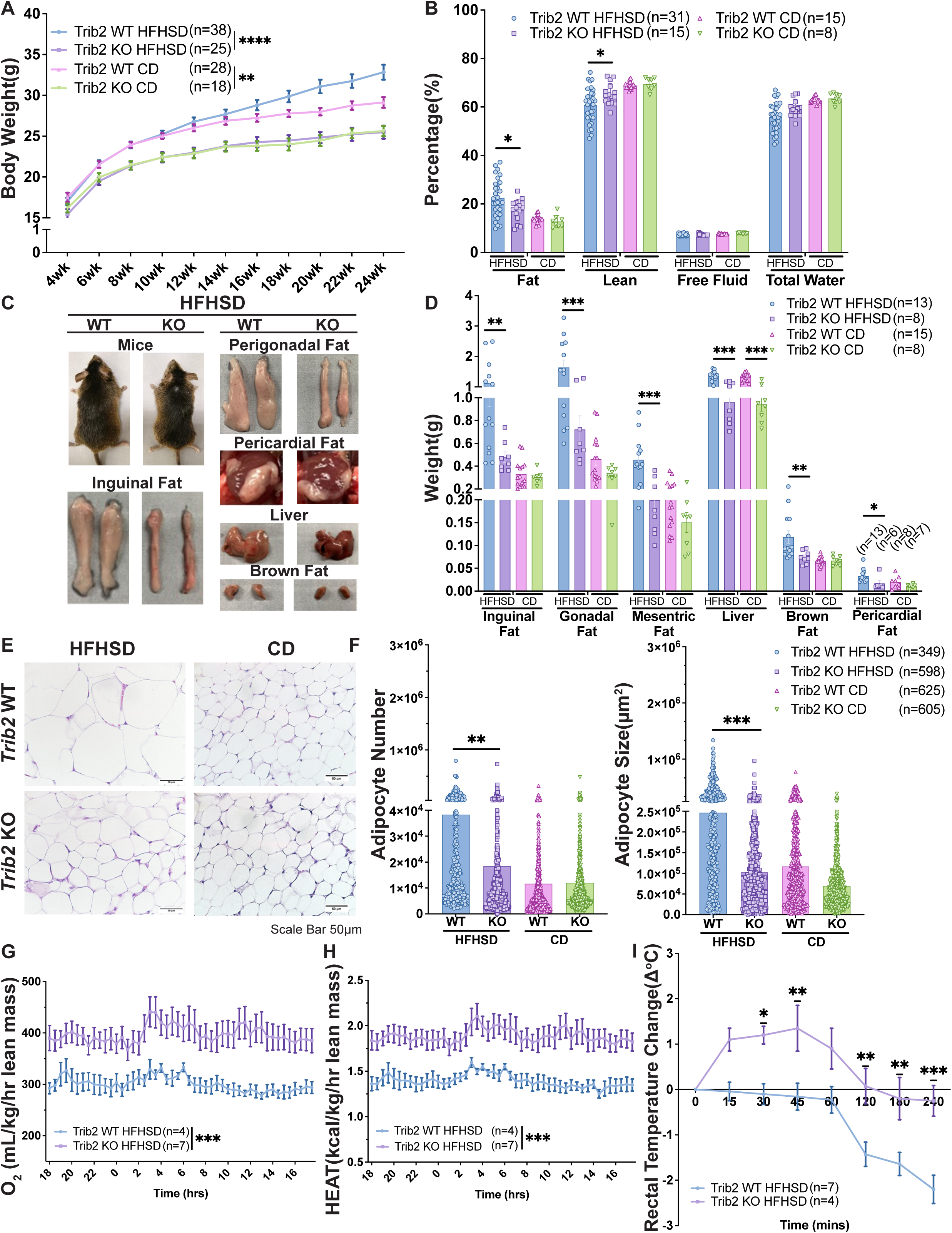
Loss of *Trib2* prevents weight gain and fat accumulation in mice while increasing energy expenditure and cold-induced thermogenesis. (A) Knockout of *Trib2* promotes weight loss in mice on a CD or a HFHSD. (B) Knockout of *Trib2* alleviates fat accumulation and preserves lean mass in mice on a HFHSD. (C&D) Knockout of *Trib2* decreased the adipose tissue and liver weights in mice fed a HFHSD. (E&F) Perigonadal adipocyte number and size were smaller in the knockout *Trib2* mice than in the WT *Trib2* mice fed a HFHSD. (G&H) *Trib2* KO mice fed a HFHSD have a high O_2_ consumption and more heat production. (I) *Trib2* KO mice fed a HFHSD maintain a higher body core temperature. Data A was analyzed using two-way ANOVA, followed by Tukey’s post hoc test for multiple comparisons. Data B, D, and F were analyzed using two-way ANOVA, followed by Šídák’s post hoc test for multiple comparisons. Data G and H were analyzed by two-way ANOVA, followed by a two-stage linear step-up procedure of Benjamini, Krieger, and Yekutiel’s multiple comparisons test. Data I was analyzed using two-way ANOVA, followed by Bonferroni’s post hoc test for multiple comparisons. All data represent the mean ± SEM. * *p* < 0.05; ** *p* < 0.01; *** *p* < 0.001; **** *p* < 0.0001.

### Loss of *Trib2* increased heat production and cold-induced thermogenesis

Obesity results from an imbalance of energy intake and energy expenditure (Hill & Commerford, 1996). The food consumption of WT and KO mice fed with HFHSD was not different (Fig EV 3A). Indirect calorimetry shows that the oxygen (O_2_) consumption and heat production (energy expenditure) were significantly enhanced in the KO mice compared to the WT mice on HFHSD (Fig. 2G&H). In addition, the KO mice maintained a higher core temperature compared to the WT HFHSD mice in the cold test (Fig. 2I). The physical activity was not different between WT and KO mice fed with HFHSD (Fig EV 3B). Hence, the loss of *Trib2* in mice resulted in increased heat production, higher energy expenditure, and thermogenesis, suggesting the reduced diet-induced weight gain of the KO mice is due to increased energy expenditure and thermogenesis.

### Deficiency of TRIB2 upregulates the UCP1 expression in brown adipose tissues and adipocytes

Non-shivering thermogenesis is maintained by brown adipose tissue (BAT) and beige adipose tissues from white adipocytes, which are characterized by the expression of uncoupling protein 1 (UCP1)(Enerbäck, 2009; Ricquier & Bouillaud, 2000). We found significantly increased UCP1 expression in BAT in the KO mice compared to WT mice under HFHSD feeding (Fig. 3A). We did not detect a difference in the white adipose tissue browning between the KO and WT mice (Fig EV 4A&B). This suggests the major energy expenditure is from the BAT. WT-1 is the mouse immortalized brown preadipocyte cell line derived from primary brown preadipocytes. The differentiation of WT1 brown preadipocytes is shown as a schematic diagram (Fig. 3B). During the differentiation of WT1 cells, we observed a decreasing level of TRIB2 expression accompanied by an increasing level of UCP1 expression (Fig. 3C, left & central panel). A negative correlation (*r*=-0.8489) between TRIB2 and UCP1 was observed from day 4 to 10 during the WT1 differentiation (Fig. 3C, right panel). TRIB2 was downregulated while UCP1 was upregulated during the differentiation. We next knock down *Trib2* in WT1 cells by shRNA. Knockdown of TRIB2 in WT1 cells upregulated UCP1 expression at both Day 4 and Day 8 after differentiation (Fig. 3D, Fig EV 5A). We further conduct a tetramethylrhodamine, methyl ester (TMRM) membrane potential functional test of the mitochondria. Knockdown of TRIB2 in WT1 cells significantly increases mitochondrial function (Fig. 3E, Fig EV 5B). These suggest that TRIB2 knockdown upregulates UCP1 expression and promotes mitochondrial function.

**Figure 3.**
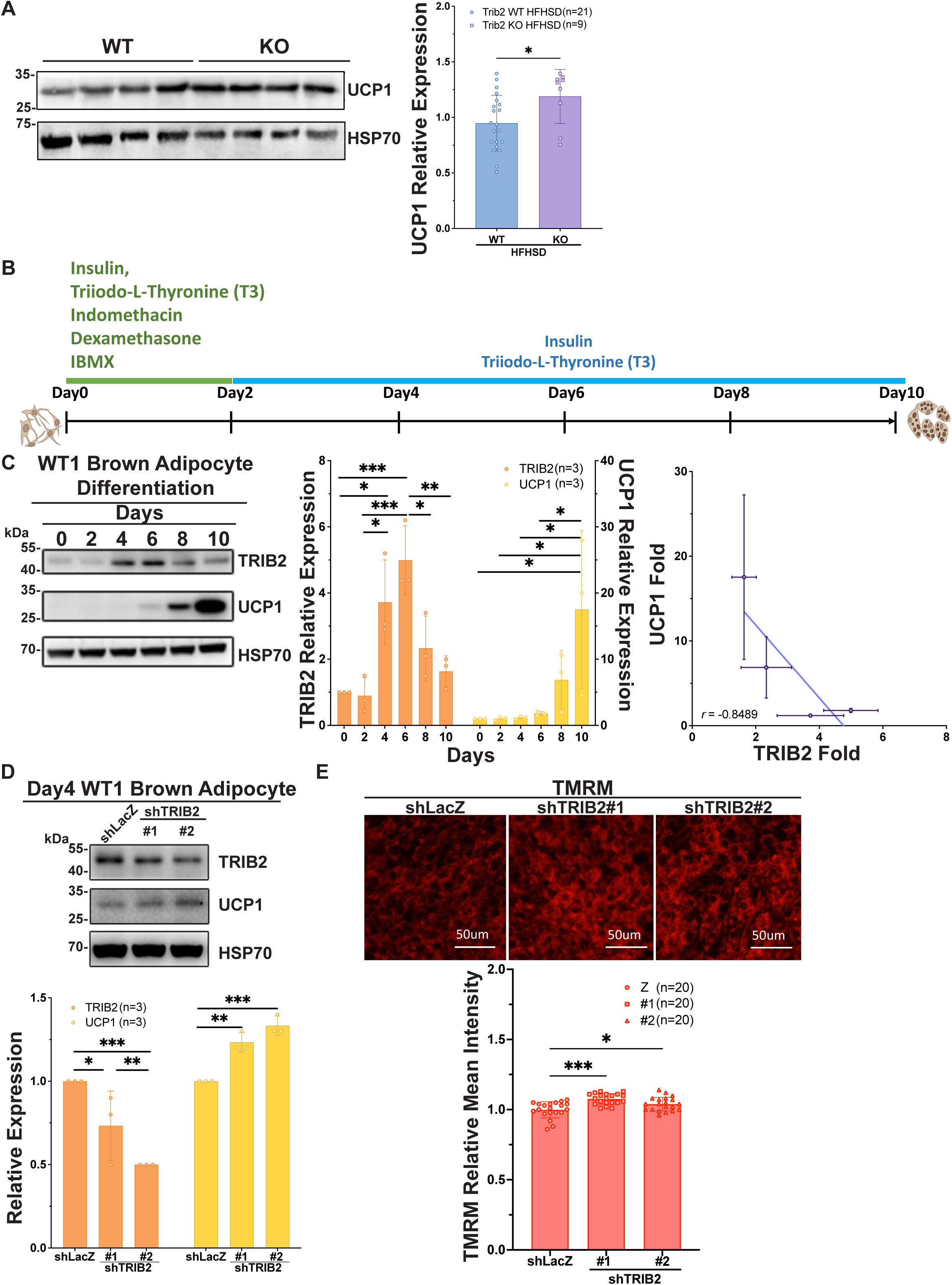
A deficiency of TRIB2 increases the UCP1 expression in brown adipose tissue and WT1 brown adipocytes. (A) The UCP1 expression increases in the *Trib2* KO mice BAT. (B) The differentiation schematic diagram of the WT1 brown preadipocyte. (C) A decreasing level of TRIB2 expression accompanied by an increasing level of UCP1 expression. The negative correlation (Pearson *r*: –0.8489) between TRIB2 and UCP1 expression during the differentiation of WT1 brown adipocytes. (D) Knockdown of TRIB2 in WT1 cells increases UCP1 expression. (E) TMRM functional test suggests that knockdown of TRIB2 in WT1 cells increases mitochondrial function. Data A, B, and E were analyzed by student unpaired t-tests. Data C correlation analysis was analyzed by Pearson’s *r* correlation. Data D was analyzed by two-way ANOVA, followed by Tukey’s post hoc test for multiple comparisons. All data represent the mean ± SD. \**p* < 0.05, \*\**p* < 0.01; \*\*\**p* < 0.001; \*\*\*\**p* < 0.0001.

### UCP1 directly binds to the TRIB2 pseudokinase domain

TRIB protein has been reported to recruit E3 ligase to degrade protein(Qi *et al*, 2006; Qiao *et al*., 2013; Xu *et al*., 2014). Knockout of TRIB2 in mice and knockdown of TRIB2 in WT1 preadipocytes increase the UCP1 expression. Hence, we hypothesized that TRIB2 directly interacts with UCP1 and leads to downregulation of the UCP1 protein level. We first transiently transfect TRIB2-His into the 293T stably expressing UCP1-GFP cells. Immunoprecipitated (IP) UCP1-GFP by GFP-Trap successfully co-precipitated TRIB2-His (Fig. 4A). Reciprocally, TRIB2-GST purified protein effectively pulls down UCP1 in BAT of WT mice (Fig. 4B). Endogenous UCP1 was able to co-precipitate endogenous TRIB2 in BAT by using the UCP1 antibody (Fig. 4C&D). We also conduct proximity ligation assay (PLA) on BAT from WT mice. *In situ* PLA signal indicated a direct interaction between endogenous TRIB2 and UCP1 (Fig. 4E). TRIB2 can be divided into three domains: the N-terminal PEST domain, the central pseudokinase domain, and the C-terminal MEK/E3 binding domain. We designed four TRIB2 truncated forms, including the full-length (TRIB2_FL) form, the C-terminal truncated form (TRIB2_NP), the N-terminal truncated form (TRIB2_PC), and the C– and N-terminal truncated ones (TRIB2_P), to elucidate which domain TRIB2 binds to UCP1 (Fig. 4F, upper panel). The TRIB2 pseudokinase (TRIB2_P) form co-precipitated with UCP1–GFP (Fig. 4F, lower panel). The results indicate that TRIB2 directly binds to UCP1, and the pseudokinase is the primary binding domain.

**Figure 4.**
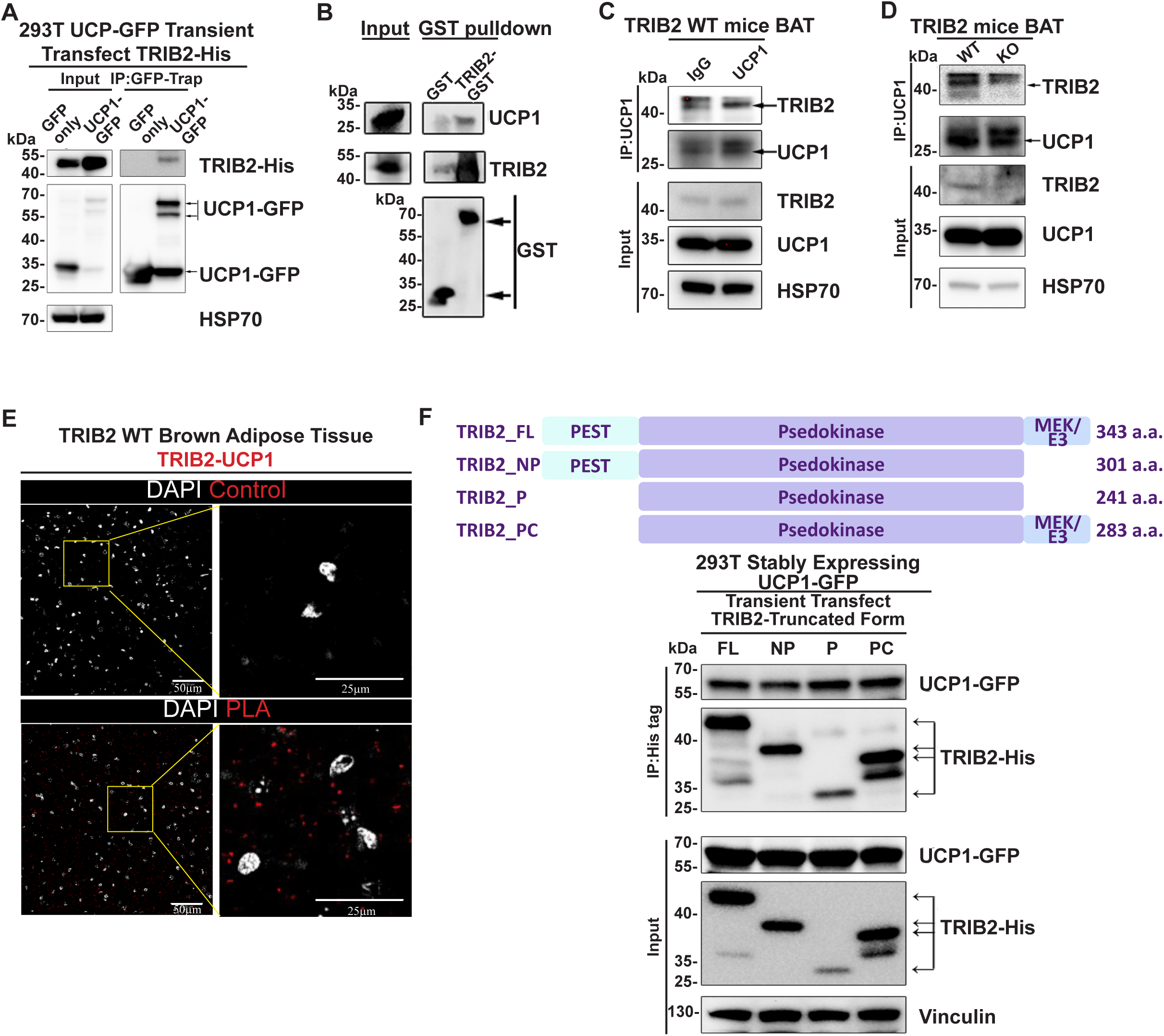
UCP1 directly binds to the TRIB2 pseudokinase domain. (A) Immunoprecipitated UCP1-GFP successfully co-immunoprecipitated TRIB2-His in 293T cells. (B) TRIB2-GST protein pull-down endogenous UCP1 in BAT. (C) Immunoprecipitated UCP1 with UCP1 antibody co-precipitated TRIB2 in BAT of WT mice. (D) Immunoprecipitated UCP1 with UCP1 antibody co-precipitated TRIB2 in BAT of WT mice, but did not co-precipitate TRIB2 in KO mice. (E) *In situ* PLA signal indicated a direct interaction between endogenous TRIB2 and UCP1 on the BAT of WT mice. (F) TRIB2-His pseudokinase truncated forms co-precipitate UCP1-GFP, suggesting pseudokinase domains can directly bind to UCP1. FL: Full length; NP: N-terminal + Pseudokinase; P: Pseudokinase only; PC: Pseudokinase + C-terminal.

### TRIB2 facilitated ubiquitination of UCP1

Given that TRIB2 is known to interact with several E3 ligases, including β-TrCP, COP1, and Smurf to degrade proteins(Qiao *et al*., 2013; Xu *et al*., 2014). Therefore, we hypothesized that TRIB2 may recruit an E3 ligase to promote UCP1 ubiquitination and its degradation. We initially transfected 293T cells stably expressing UCP1-GFP with varying doses of Ubiquitin-HA. Immunoprecipitation UCP1-GFP revealed a dose-dependent increase in its ubiquitination (Fig. 5A). This suggests UCP1 can be ubiquitinated in cells. We co-transfected 400ng Ubiquitin-HA and varying doses of TRIB2 into 293T cells. 2µg of TRIB2-His combined with 400ng of Ubiquitin-HA results in downregulated UCP1 expression (Fig. 5B). Subsequently, we co-transfected 400 ng of Ubiquitin-HA with varying doses of TRIB2-His into 293T cells and treated them with MG132 for 16 hours. Immunoprecipitation of UCP1-GFP revealed increased ubiquitination of UCP1 in the presence of TRIB2 (Fig. 5C). We further validated ubiquitination of UCP1 in the knockdown of TRIB2 in WT1 cells. TRIB2 knockdown resulted in significantly decreased UCP1 ubiquitination, along with a significant upregulation of UCP1 expression (Fig. 5D). These findings strongly suggest that TRIB2 likely facilitates the ubiquitination of UCP1.

**Figure 5.**
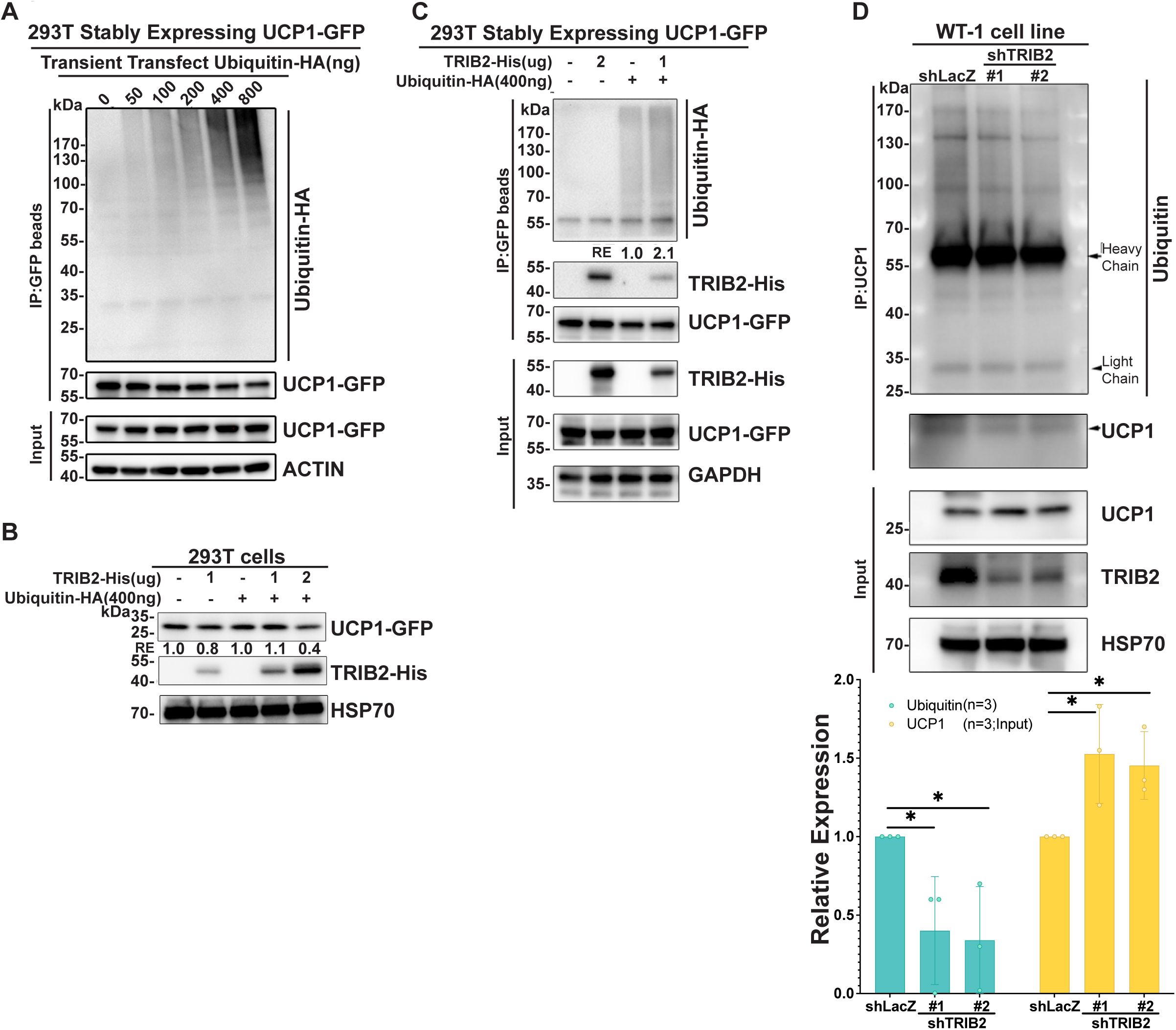
TRIB2 facilitated ubiquitination of UCP1. (A) A dose-dependent increase in the ubiquitination of UCP1. (B) Co-expressing TRIB2-His and Ubiquitin-HA in 293T cells decreases UCP1 expression. (C) TRIB2 enhances ubiquitination of UCP1. (D) Knockdown of MYCBP2 decreases ubiquitination of UCP1 in WT1 cells. Data C was analyzed by student unpaired t-tests. All data represent the mean ± SD. \**p* < 0.05, \*\**p* < 0.01; \*\*\**p* < 0.001; \*\*\*\**p* < 0.0001.

### MYCBP2 directly binds UCP1 and, together with TRIB2, facilitates its degradation

We next sought to identify potential E3 ligases involved in UCP1 ubiquitination. Given that β-TrCP, COP1, and Smurf1 are known to interact with TRIB2, we first tested whether UCP1 co-precipitates with these ligases; However, no interaction was detected, suggesting that these TRIB2-associated E3 ligases are not responsible for UCP1 degradation (Fig. 6A). To systemically identify the E3 ligase interacting with UCP1, we performed immunoprecipitation of UCP1 followed by LC-MS/MS analysis (Fig. 6B). MYCBP2, an E3 ligase, was identified among the 129 identified proteins (Table1). MYCBP2 is known to exist at approximately 510 kDa in the embryonic state and 350 kDa in the adult state(Santos *et al*, 2006). We co-immunoprecipitated in adult mouse BAT, where UCP1 co-precipitated with MYCBP2 at 350 kDa (Fig. 6C). In 293T cells transiently transfected with MYCBP2-Myc and UCP1-GFP, UCP1-GFP co-precipitation confirmed a direct interaction of MYCBP2 at 510kDa (Fig. 6D). Furthermore, *in situ* PLA performed on BAT specimens demonstrated a direct interaction between endogenous MYCBP2 and UCP1 (Fig. 6E). However, the interaction between MYCBP2 and UCP1 did not find on KO BAT, suggesting this interaction may through the TRIB2 (Fig. 6F). Varying doses of MYCBP2 were transfected into 293T cells, resulting in a dose-dependent downregulation of UCP1 expression (Fig. 6G). Next, we knocked down MYCBP2 in WT1 cells using shRNA. MYCBP2 knockdown efficiently upregulates UCP1 expression, and a negative correlation (*r*=-0.9981) between MYCBP2 and UCP1 was observed (Fig. 6H). We further performed a TMRM assay to assess mitochondrial function, which revealed that MYCBP2 knockdown effectively enhanced mitochondrial membrane potential (Fig. 6I).

**Figure 6.**
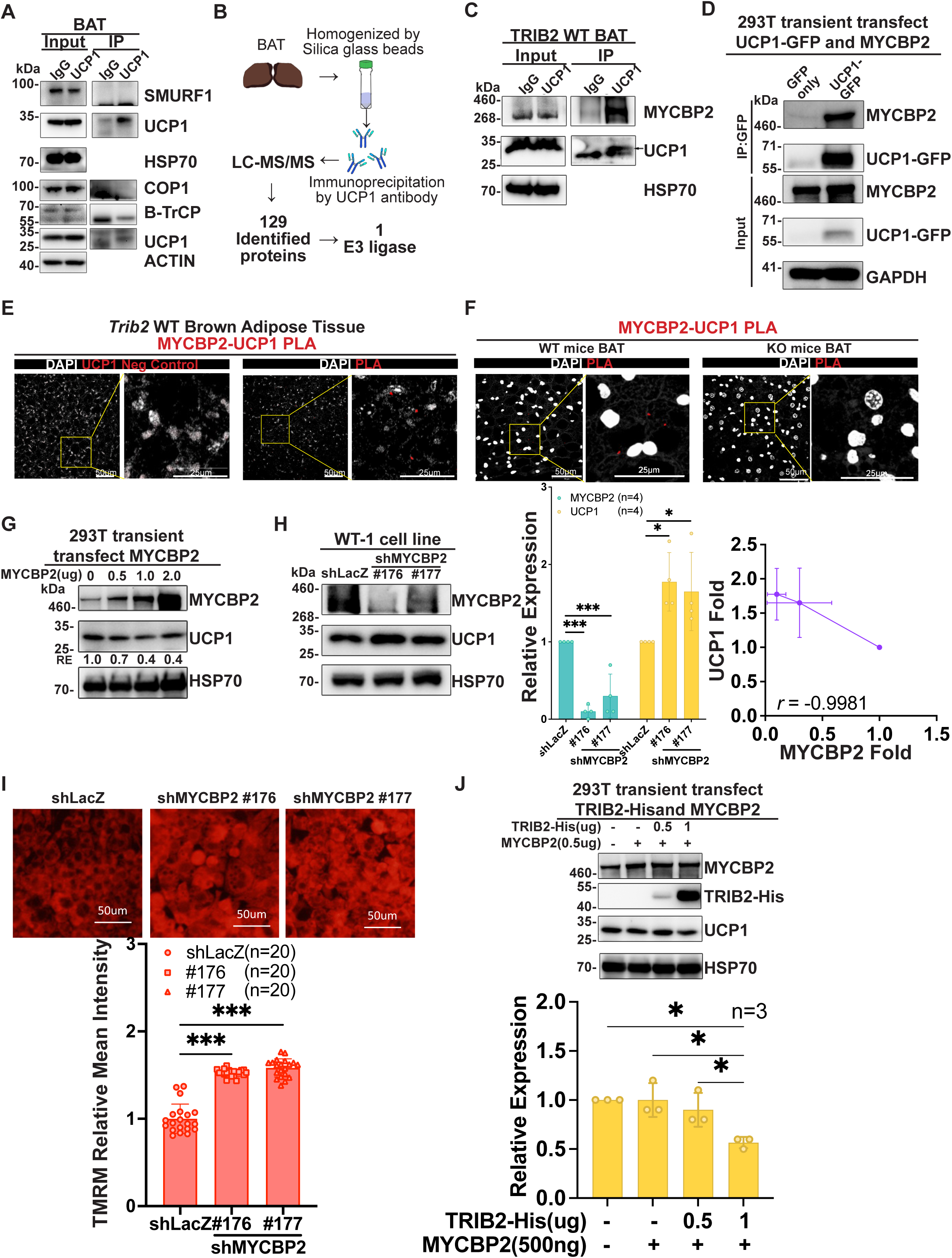
TRIB2 and MYCBP2 cooperatively promote UCP1 degradation. (A)Immunoprecipitated UCP1 with UCP1 antibody did not co-precipitate the TRIB2 reported interacting E3 ligase. (B) The scheme for identifying UCP1-interacting proteins by LC-MS/MS. (C) Immunoprecipitated UCP1 with UCP1 antibody co-precipitates MYCBP2 in BAT of WT mice. (D) Immunoprecipitated UCP1-GFP co-precipitates MYCBP2 in 293T cells. (E&F) *In situ* PLA signal indicated a direct interaction between endogenous UCP1 and MYCBP2 on the BAT of WT mice, but not on the BAT of KO mice. (G) Overexpressing MYCBP2 in 293T decreases UCP1 expression. (H) Knockdown of MYCBP2 increases UCP1 expression in WT1 cells. The negative correlation (Pearson *r*: –0.9981) between MYCBP2 and UCP1. (I) TMRM functional test suggests that knockdown of MYCBP2 in WT1 cells increases mitochondrial function. (J) Expressing TRIB2 enhances MYCBP2 to decrease UCP1 expression in 293T cells. Data H was analyzed by two-way ANOVA, followed by Tukey’s post hoc for multiple comparisons and Pearson’s *r* correlation. Data I was analyzed by student unpaired t-tests. Data J was analyzed by one-way ANOVA, followed by Tukey’s post hoc for multiple comparisons. All data represent the mean ± SD. \**p* < 0.05, \*\**p* < 0.01; \*\*\**p* < 0.001; \*\*\*\**p* < 0.0001.

Additionally, in contrast to Fig. 5B, co-expression of 1 µg TRIB2 and 0.5 µg MYCBP2 effectively suppressed UCP1 expression in 293T cells (Fig.6J). These findings suggest that TRIB2 and MYCBP2 cooperate to regulate UCP1 expression. Taken together, TRIB2 acts as a scaffold protein to recruit the E3 ligase, leading to the ubiquitination and degradation of UCP1. Loss of TRIB2 increases UCP1 expression, thereby enhancing thermogenesis and contributing to altered fat distribution and improved metabolic health.

## Discussion

In this study, we identified a previously unrecognized regulatory mechanism of UCP1 through E3 ligase–mediated ubiquitination by TRIB2. We demonstrated that *Trib2* deficiency prevented diet-induced obesity, reduced fat mass, increased lean mass, ameliorated hepatic steatosis, and enhanced glucose tolerance and insulin sensitivity through enhanced thermogenesis. At the molecular level, TRIB2 negatively regulates UCP1 protein abundance via proteasomal degradation. Mechanistically, TRIB2 directly interacts with UCP1 through its pseudokinase domain and facilitates its ubiquitination by recruiting the E3 ligase MYCBP2 (Fig. 7). These findings reveal ubiquitination and proteasomal degradation as a new post-translational regulator of UCP1-mediated thermogenesis and present a promising therapeutic approach, analogous to proteolysis-targeting chimera (PROTAC) strategies.

**Figure 7.**
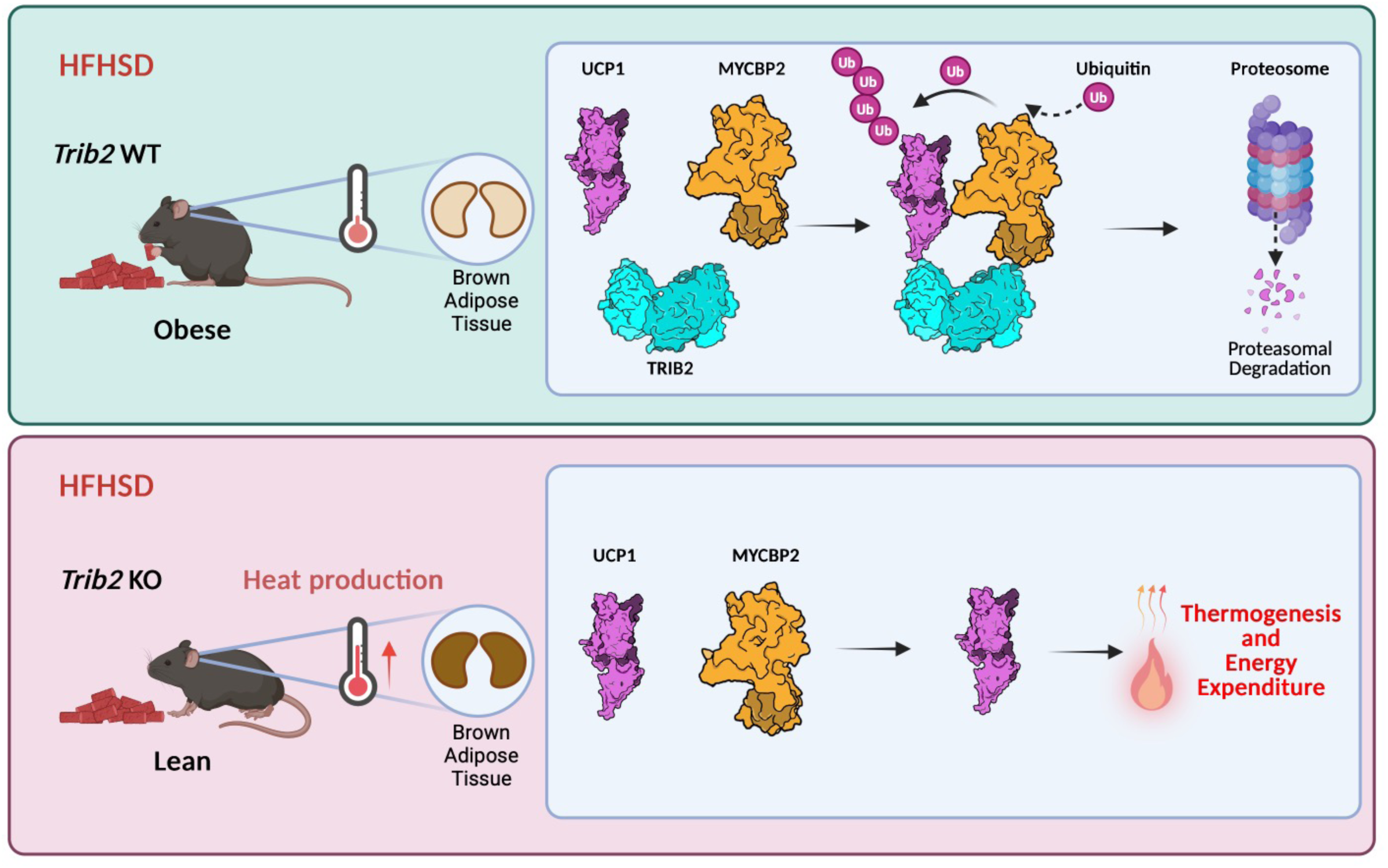
Graphic abstract. *Trib2* knockout mice displayed elevated UCP1 expression in brown adipose tissue, leading to enhanced thermogenesis and increased energy expenditure. Mechanistically, TRIB2 binds directly to UCP1 via its pseudokinase domain and promotes its ubiquitination by recruiting the E3 ligase MYCBP2.

Previous studies have suggested a role for TRIB2 in fat distribution and adiposity. For example, Fox et al. identified a SNP near TRIB2 (rs10198628, MAF 0.49, p = 2.7 × 10⁻⁸) that was significantly associated with pericardial fat (p = 5.4 × 10⁻¹⁴) in a meta-analysis of GWAS involving 5,487 individuals of European ancestry from the Framingham Heart Study and the Multi-Ethnic Study of Atherosclerosis (Fox *et al*., 2012). A similar result was reported by Chu et al., who confirmed the association of rs10198628 with pericardial fat (P < 5 × 10⁻⁸)(Chu *et al*., 2017). In East Asian populations, Nakayama et al. identified a different SNP in the 3’UTR of TRIB2 (rs1057001), which was associated with visceral fat area (VFA; p = 2.7 × 10⁻⁶), waist circumference (WC; p = 9.0 × 10⁻⁶), and waist-to-hip ratio (WHR; p = 0.017)(Nakayama *et al*., 2013) in a candidate genetic association study. Similar associations were reported by Wang et al., where rs1057001 was significantly associated with WC in two statistical models (Wang *et al*., 2016). Furthermore, Nakayama and Iwamoto et al. demonstrated that rs1057001 correlates with higher expression levels of thermogenic genes in both subcutaneous and visceral human adipose tissues(Nakayama & Iwamoto, 2017). The A allele of rs1057001 was uncommon in sub-Saharan Africans but highly prevalent in East Asians and Siberians. Consistently, we demonstrated that the A allele of rs1057001 caused instability of TRIB2 RNA. Collectively, these studies suggest that TRIB2 influences fat accumulation, thermogenesis, and correlates with cold adaptation during human migration.

In addition to a variety of factors, including cold exposure, diet, hormones, purine-nucleotide, and certain drugs such as peroxisome proliferator-activated receptor gamma (PPARγ) agonists and β3-adrenergic agonists, as well as transactional or epigenetic control, other modifications of UCP1 have been identified. Evanna L et al. elegantly showed that reversible covalent sulfenyl modification of UCP1 by ROS regulates thermogenesis(Mills *et al*, 2022). Wang, et al. revealed that reversible lysine succinylation and malonylation of UCP1 decreased its stability and activity and impaired mitochondrial metabolism(Wang *et al*, 2019). Johnson, et al. found that phosphatidylethanolamine (PE) is an important component of the mitochondrial inner membrane that modulates UCP1-dependent proton conductance and promotes thermogenesis(Johnson *et al*, 2023). In addition, the polyadenylation of the 3’UTR of UCP1 as well as lysine acetylation also influence UCP1 stability and function, respectively(Lu *et al*, 2020). Here, we identified another new regulatory pathway of UCP1 by E3 ligase-mediated protein degradation, which may be related to environmental temperature during human migration.

The Tribbles (TRIB) pseudokinase family consisted of TRIB1-3(Shrestha *et al*., 2020). They evolved unique features distinguishing them from all other protein kinases. The atypical pseudokinase domain retains a regulated binding platform for substrates, which are subsequently ubiquitinated by context-specific E3 ligases(Qiao *et al*., 2013; Wang *et al*., 2013). In two large independent GWAS, genetic variants of *TRIB1* are associated with serum TG levels(Kathiresan *et al*, 2008; Willer *et al*, 2008). Hepatic-specific overexpression of *Trib1* reduced levels of serum TG and cholesterol by reducing VLDL (very low-density lipoproteins); conversely, *Trib1* knockout mice showed elevated levels of serum TG and cholesterol due to increased VLDL production. Another structural and mutagenesis study clearly showed that the pseudokinase domain TRIB1 is able to interact with C/EBP proteins and regulate their degradation by recruiting E3-ligase constitutive photomorphogenic protein 1 (COP1)(Niespolo *et al*, 2020). Transgenic mice expressing TRIB3 in adipose tissue are protected from diet-induced obesity due to increased lipolysis by triggering the ubiquitination-mediated degradation of acetyl–coenzyme A carboxylase (ACC), the rate-limiting enzyme in fatty acid synthesis through E3 ligase COP1(Qi *et al*., 2006). Collectively, the TRIB family represents a unique family regulating lipid and energy metabolism by ubiquitination-mediated degradation of key enzymes.

In consistent with our findings, Wang et al. demonstrated that TRIB2 deficiency ameliorates high-fat diet (HFD)-induced hepatic insulin resistance, inflammation, and endoplasmic reticulum stress in mice(Wang *et al*., 2024). Mechanistically, they found TRIB2 promotes hepatic insulin resistance by interacting with the AMPK β subunit (PRKAB), disrupting the AMPK complex via its pseudokinase domain. Since AMPK activation suppresses lipogenesis by inactivating Acetyl-CoA Carboxylase (ACC), TRIB2-mediated inhibition of AMPK may drive hepatic lipid accumulation and insulin resistance. Their study highlights TRIB2’s scaffolding function in the liver, where it modulates AMPK signaling. In contrast, our study identifies a novel role for TRIB2 in brown adipose tissue (BAT), where it binds UCP1 and recruits the E3 ligase MYCBP2 to promote UCP1 ubiquitination and degradation. This action suppresses thermogenesis and contributes to obesity. Together, these findings suggest that TRIB2 functions as a multifaceted regulator of energy homeostasis and metabolism across different tissues. Importantly, TRIB2 has emerged as a target of tyrosine kinase inhibitors—originally developed to inhibit HER2 and EGFR—such as afatinib, neratinib, osimertinib (Foulkes *et al*, 2018), and daclatasvir(Monga *et al*, 2023). These drugs have been recently found to destabilize TRIB2, suggesting a potential therapeutic strategy.

The further investigation could focus on the following aspects. First, we did not employ tissue-specific knockout models such as *Pdgfra*-*cre* or *Ucp1*-*cre* mice(Reverte-Salisa *et al*, 2024), nor did we utilize tissue-targeted delivery systems like AAV8(Tsuji *et al*, 2023) or Rec2(Huang *et al*, 2019) viruses, which would allow for more precise assessment of TRIB2 function specifically in brown adipose tissue. Second, although our results support enhanced thermogenesis as measured by indirect calorimetry, we did not perform *in vivo* metabolic flux analyses, such as positron emission tomography (PET) using ^18^F-fluorodeoxyglucose (^18^F-FDG)(Haley *et al*, 2023; Seki *et al*., 2022), to directly assess the uptake of ^18^F-FDG in BAT. Third, while we observed improved glucose homeostasis in *Trib2*-deficient mice, we did not perform a euglycemic hyperinsulinemic clamp for assessment of insulin sensitivity, which would have provided a more quantitative evaluation of insulin sensitivity. Fourth, we did not investigate the phenotypic, metabolic, or physiological effects of small-molecule kinase inhibitors that target TRIB2, such as afatinib, and the potential off-target effects of these inhibitors—originally developed to target EGFR/HER2—must be carefully evaluated in the metabolic context. Lastly, the long-term effects and safety of modulating the TRIB2–UCP1 axis remain unknown and warrant further investigation. Addressing these limitations in future studies will provide a more comprehensive understanding of TRIB2’s role in energy metabolism and therapeutic feasibility.

Taken together, we identified a new regulatory mechanism of UCP1 level through ubiquitination-mediated degradation by TRIB2. Human genetic studies strongly support that genetic variants of TRIB2 are associated with thermogenesis, fat accumulation, and human migration related to ambient temperature. With the identification of more small-molecule TRIB2-targeting compounds with minimal off-target effects, targeting the TRIB2–UCP1 axis may be a new therapeutic strategy analogous to PROTAC techniques.

## Materials and Methods

### Animal model

The *Trib2* global knockout mice were created by deleting exons 1 and 2, resulting in the absence of TRIB2 protein production. A neomycin resistance cassette (Neo) was integrated into exons 1 and 2 to generate mouse embryonic stem cells lacking *Trib2*. Initially, homologous recombination replaced the *Trib2* gene with a *lacZ* reporter gene and a *lox*P-flanked neomycin resistance gene (Neo^r^) cassette. Subsequently, *Cre-LoxP* recombination facilitated the removal of the Neo cassette by acting on the flanking *loxP* sites. The mice were bred and maintained in accordance with the guidelines of the Animal Center at the National Taiwan University College of Medicine, which is accredited by AAALAC (Association for Assessment and Accreditation of Laboratory Animal Care International). They were housed under a 12-hour light-dark cycle at 23°C and provided *ad libitum* access to a regular chow diet (Lab Diet). Ethical approval for the study was obtained from the Institutional Animal Care and Use Committee of the Medical College of National Taiwan University (approval numbers: 20170237 and 20210005).

### Body composition

Body composition was determined by Skyscan 1076 in vivo micro-CT scanner (Skyscan, Bruker) with the average of three repeated measurements to calculate body fat, lean mass, free fluid, and total water. Mice with both a regular chow diet or HFHSD were measured at 28∼30 weeks old.

### Glucose and insulin tolerance test

For the glucose tolerance test, the intraperitoneal glucose tolerance test (i.p.GTT) was evaluated with a 6-hour fasting period previously. 1mg/g glucose (20% Vitagen, Taiwan Biotech) diluted with 0.9% isotonic sodium chloride solution (Taiwan Biotech) was given by i.p. and measured tail blood glucose with ACCU-CHEK Performa glucometer (Roche) at 0, 15, 30, 45, 60, 90, and 120 minutes after injection. For the insulin tolerance test, the intraperitoneal insulin tolerance test (i.p.ITT) was evaluated with a 4-hour fasting period previously. 0.1 IU/g of insulin (100 IU/mL insulin dilute solution similar to i.p.GTT, Humulin R, Eli Lilly) was also given by i.p., and the tail blood glucose was measured at 0, 15, 30, 45, 60, 90, 120, and 180 minutes after injection. Area under the curve (AUC) and inverse AUC were calculated in response to i.p.GTT and i.p.ITT, respectively.

### Energy expenditure

Indirect calorimetry and physical activity measurements were performed using the CLAMS-HC system (Comprehensive Lab Animal Monitoring System for HOME CAGES, Columbus Instruments) at Taiwan Mouse Clinic. The input and output concentration of the chambers for oxygen consumption and carbon dioxide production (as VO_2_ and VCO_2_) were monitored, and the respiratory quotient (the ratio of CO_2_ to O_2_) was calculated. Further, the heat released from the body’s calorific value was computed by gas exchange, accounting for the energy released during the metabolic process. The running wheel was also placed in the home cage for physical activity measurement. With the body weight difference between wild-type and knockout mice, instead of taking the exact weight of mice for the experiment, we took the mice to meet the average weight in each one and adjusted metabolic rate (MR) by body lean mass normalization. The normalization method of mass-adjusted MR was based on Muller, T. D., et al.’s study(Müller *et al*, 2021). Before the MR measurement, mice were fed with HFHSD for 1 week, measured body composition, and accommodated single-housed in the CLAMS system for 3 days. The mean daily MR of each individual was continuously monitored for 3 days. Quantifying dietary intake, HFHSD were fed ad libitum and weighed daily food consumption for 7 continuous days. Food intake was measured with housing in group cages with the same genotype, providing for individual mouse average intake.

### Cold-induced thermogenesis (CIT)

For thermogenic activation, mice completed an hour of cold exposure one week before the thermogenesis test. Mice were tested at 37-41 weeks old with HSFSD from 4 weeks old. Fasting 4 hours before the test, CIT was performed on single-housed without bedding, with drinking, re-feeding HFHSD, and in 4℃ chambers. Body temperature was measured in conscious mice using a rectal probe TH-5 Thermalert Clinical Monitoring Thermometer (Physitemp). The time course measurement was set as the first hour with four intervals and measured 1 hour each, respectively, within a total of 4 hours.

### Histology and Immunohistochemistry

For histopathological examination, perigonadal fat and liver were incubated and fixed in 4% formalin, embedded in paraffin blocks, and sectioned at a thickness of 20µm for hematoxylin and eosin (H&E) stain. Following the conventional procedure, deparaffinization, hydration, hematoxylin, and eosin staining. Examining histological differences was captured under light microscopy Olympus BX51 microscope combined with Olympus DP72 camera and CellSens Standard 1.14 software (Olympus) or MIRAX SCAN (Carl Zeiss). Immunohistochemistry (IHC) was performed on a BOND-MAX fully automated IHC staining system (Leica). 20 µm paraffin sections were baked, deparaffinized, and stained in this instrument. Slides for mouse staining were antigen-retrieved at pH 9, 95 °C, 30 min. F4/80 antibody (Abcam; 1:100) and UCP1 antibody (#GTX10983; 1:200) were incubated for 60 minutes at room temperature. Slides were stained with a DAB detection kit (Bond refine systems, Leica) for 5 min and counterstained with hematoxylin (Bond refine systems, Leica) for 7 min. All slides were mounted with a coverlid and scanned at 40x magnification by Aperio GT450 (Leica).

### Quantification of adipose cell size and number

The sections were counted under the 400x magnification and captured under light microscopy, Leica DM IL LED Inverted Microscope (11521257/343358). For cell size, follow the steps, using ImageJ with segmented particle analysis, selecting the adipocytes area with the wand tools, recording the measurements, and converting pixels to µm. For cell number, fat pad mass is divided by 0.9 for fat density and then divided by cell size (cm^3^).

### Blood biochemistry analysis

For hepatic TG content, liver tissue TG extraction and quantification were performed using the TG assay kit (Wako) and normalized by the tissue weight. Blood was collected from a fasting specimen and separated by centrifugation at 800 g at 4℃ for 5 minutes for serum lipid profiling. The level of Triglyceride (TG) was measured using Cobas c 111 analyzers (Roche). The level of adiponectin (1:10’000) and leptin (1:5) levels were measured using DuoSet ELISA Development Kits.

### Mitochondrial membrane potential assay

Tetramethyl rhodamine methyl ester (TMRM) (ThermoFisher) was a cell membrane-permeable cationic fluorescent dye for a polarized cell that will have higher dye uptake to quantify the change of mitochondrial membrane potential. Followed the manufacturer’s instructions. Primary brown adipocytes and WT-1 cells were prepared for assay. Cells were treated with 50nM TMRM reagent in serum-free culture medium, incubated for 30 minutes, washed twice with PBS, and replaced with HBSS-HEPES imaging buffer by supplementing 20mM HEPES (Bionovas) in Hank’s Balanced Salt Solution. Consequently, the fluorescence intensity of living cells was captured under an inverted fluorescence microscope Olympus IX71, combined with Olympus DP70 camera and CellSens Standard 1.14 software (Olympus). Cell fluorescence intensity was measured and quantified with FIJI.

### Vector Cloning

Full-length mouse *Trib2* DNA cloning on pGEX-4-T1 vector. Mouse *Trib2* cDNA and pGEX-4-T1 were digested by BamHI (#R0136S) and XhoI (#R0146S) enzymes using NEB buffer 3.1 (#B7203). Full-length human *TRIB2* DNA cloning on pcDNA3.1/Myc-His (+) A (#V80020) vector. Human *TRIB2* and pcDNA3.1/Myc-His (+) A were digested by HindIII-HF (#R3014S) and XhoI (#R0146S) enzymes using NEB buffer rCutsmart (NB6004S). UCP1 Human tagged ORF clone (#RC218901) was purchased from OriGene Technologies and subcloned into EcoRI (#R3101S) and XmaI (#R0180S) sites of pLV-C-GFPSpark Lentivirus vector (#LVCV-35). pCMV-Pam-FL was a gift from Vijaya Ramesh^89^ (Addgene plasmid # 42570; http://n2t.net/addgene:42570; RRID: Addgene_42570). All cDNA and vectors were ligated by T4 ligase (#M0202S) using 10x T4 ligase Buffer (#B0202S). *TRIB2* 3’UTR was cloned from genomic DNA (#D1234999), ligated into Dual-Glo Luciferase Vector (#E2940) as Hap1 vector, and further site-directed mutated to generate Hap2 vector. For the knockdown of *Trib2* in WT-1 cell lines, the negative control is pLKO.1-puro-sh*LacZ* (Cat. #TRCN0000072223) and pLKO.1-puro-sh*Trib2* vectors (Cat. #TRCN0000024216 and TRCN0000024218) purchase from National RNAi Core Facility.

### Lentivirus and Infection

Knockdown *Trib2* by lentiviral transduction to introduce short hairpin RNA (shRNA). Co-transfect lentiviral construct, packaging plasmid pSAX2, and envelope plasmid pMD2G in HEK-293T cells (293T cells; #CRL-3216) to produce lentivirus. Collect the lentivirus-containing supernatant after 48 hours of transfection. WT-1 cell lines were infected with lentivirus and selected with 20µg/ml puromycin. HEK-293T cells were infected with lentivirus to overexpress UCP1 (293T_UCP1-GFP) and sorted for GFP-positive cells by flow cytometry (FACS).

### Cell Culture

WT-1 mouse brown preadipocyte (#SCC255) was purchased from Merck Ltd, and HEK-293T (Cat. #CRL-3216) was obtained from ATCC. WT-1 preadipocytes and HEK-293T cells were cultured in Complete DMEM Medium (Dulbecco’s Modified Eagle Medium (#11965092) supplemented with 10% FBS, 1% Sodium Pyruvate, 1% Non-Essential Amino Acids,1% L-Glutamine, and 0.2% Antibiotic-antimycotic). WT-1 cells should be passaged around 80-90% confluence. For WT-1, preadipocytes differentiate to mature brown adipocytes, adding Induction Medium (20nM Insulin, 1nm Triiodo-L-Thyronine (T3), 0.125mM Indomethacin, 5µM Dexamethasone, and 0.5mM IBMX in Completed DMEM medium) for two days and substituted to Differentiation Medium (20nM Insulin and 1nm Triiodo-L-Thyronine (T3) in Completed DMEM medium) every other day.

### Lipid Droplet Staining

Accumulation of lipid droplets was quantified by Oil red O (#01391) staining. Removed the medium of WT1 cells and washed with PBS. Then, cells were fixed with 4% paraformaldehyde (#158127) for 10 minutes at room temperature and washed with PBS. Added diluted Oil red O 500µl in each well and incubated for 30 minutes at room temperature. Remove the Oil red O and carefully wash it with tap water for 30 minutes at room temperature. Keep the samples in the PBS and take images to record. After recording, 100% isopropanol was used to extract the oil red O and measure it at 510nm.

### Duolink *In Situ* Proximity Ligation Assay

Duolink *In Situ* PLA (#SI-DUO92101) was obtained from MERCK Ltd and used to detect protein-protein interactions following the manufacturer’s protocol. Briefly, brown adipose tissues, antigen retrieval (pH6 ER1, 30 minutes) was performed, followed by a 30-minute incubation at 37℃ for blocking with Duolink blocking solution. BATs were incubated with primary antibodies (anti-TRIB2, #GTX83495; anti-UCP1, #GTX112784 and #sc-293418; anti-MYCBP2, #ab86078; dilution factor 1:100) in Duolink antibody diluent overnight at 4℃. Next, BATs and WT-1 cells were washed with Duolink Buffer A and hybridized with Duolink *In Situ* PLA Probe Anti-Rabbit PLUS (#DUO92002)/ Anti-Mouse MINUS (#DUO92004) species-specific probes for 1 hour at 37℃. After ligation and amplification using Duolink *In Situ* Detection Reagents Red (#DUO92008), BATs and cells were washed with Duolink Buffer B and mounted with Duolink *In Situ* Mounting Medium with DAPI (#DUO82040). The signal was captured using a high-speed confocal microscope (Leica SP8X Confocal Super-resolution Microscope). DAPI-labeled signals were excited at 405 nm, and the emitted signals were collected through a 430–480 nm bandpass filter. For the Duolink *In Situ* Detection Reagents Red, excitation occurred at 594 nm, and the fluorescence was collected using a 600–680 nm bandpass filter. Fluorescence signals from the labelled structures are pseudo-coloured.

### Protein Purification

Transform pGEX-4T-3-GST and pGEX-4T-3-m*Trib2*-GST separately into BL21 competent cells (#FYE207-40VL). Inoculate a single colony and incubate it in 10 ml LB broth with ampicillin overnight at 37℃. Inoculate 10 ml of the overnight culture the following day into 100 ml LB broth with ampicillin. Allow the cells to incubate until the OD600 reaches 0.6∼0.8 (approximately 60 minutes). Induce m*Trib2*-GST protein expression by adding 0.1 mM Isopropyl-β-D-1-thiogalactopyranoside (IPTG) and continue incubation for 4 hours. Harvest the cells by centrifugation at 4000g for 15 minutes. Discard the supernatant and resuspend the pellet in buffer R (50mM Tris-HCl, pH 8.0; 100mM NaCl; 1mM EDTA; 1mM DTT; 1mM PMSF; 0.2% Triton X-100; Protease inhibitor). Disrupt the cells using a microfluidizer. Harvest the supernatant by centrifugation at 4000g for 15 minutes. Pre-balance the GST beads (#25237) with buffer R and pass the supernatant through the GST column or incubate the supernatant with GST beads overnight. Wash the beads with at least 10 column volumes (CV) of buffer R. Assess the protein’s purity by running an SDS gel and staining it with Instant Blue (#ab119211). Store the protein of the beads at – 20°C for further use.

### Pull-down Assay

For pulldown assay, take the BATs from *Trib2* WT or KO mouse and homogenize the BAT in RIPA buffer (20 mM Tris-HCl, pH 7.5, 150 mM NaCl, 1 mM EDTA, 1% NP-40, 0.1% SDS, and 1% sodium deoxycholate) by using Silica glass beads. Next, centrifuge at 12000g for 10 minutes at 4℃ and harvest the lysate. Quantify the protein concentration by using the Bradford method. Proximity 2 mg of protein from BAT, set aside 40µg for input, and aliquot the remaining lysate into two portions for pulldown assays. Add purified 5µg GST-protein beads and 5µg mTRIB2-GST protein beads into the remaining lysate, then incubate overnight at 4℃. Centrifuge at 12000g to get the beads and wash with RIPA buffer at least 3 times. Add 2x Sample Buffer and incubate at 95℃ for 10 mins. The samples were ready for Western Blot.

### Immunoblotting

Briefly, cells and tissues were sonicated or homogenized in RIPA buffer. Collect the supernatant after centrifugation at 12000g for 10 mins at 4℃. Protein will be quantified by Bradford assay and then mixed with Laemmli Sample buffer (50mM Tris pH 6.8, 10% Glycerol, 1% SDS, 0.02% Bromophenol blue, and 20% 2-mercaptoethanol). For the ubiquitination collection experiment, 10µM MG132 was treated in the cell medium for 16 hours before the collection. After 16 hours of treatment, collect cells by RIPA with 10µM MG132 and 10mM NEM. For gel electrophoresis, 20µg of protein was loaded in a 4-12% Gradient Gel, separated and transferred to PVDF membrane (#IPVH00010) by Bio-Rad transblot system in transfer buffer (#IB3352; 50mM Tris, 40mM Glycine, 0.04%SDS and 20% Methanol). For MYCBP2 protein immunoblotting, 20µg of protein was loaded in a 3-8% Gradient Gel (#NP0321BOX), run with 150V for an hour, and transferred to PVDF membrane with 33V for 18 hours. After blocking with skimmed milk at room temperature for an hour, the membrane was incubated with primary antibody (anti-TRIB2(#13533S; 1:1000); anti-UCP1(#GTX10983; 1:2000); anti-MYCBP2 (#ab86078; 1:1000); anti-Ubiquitin (#GTX128826; 1:1000); anti-GFP (#GTX113617; 1:10000); anti-His (#HRP-66005; 1:10000); anti-GST(homemade;1:5000); anti-SMURF1(sc-100616; 1:1000); anti-COP1 (#ab56400; 1:1000); anti β-TrCP (#4394S; 1:1000); anti-HSP70 (#GTX61394; 1:1000); anti-Vinculin (#GTX113294;1:10000); anti-actin (#GTX629630; 1:10000); anti-GAPDH (#GTX100118; 1:10000)) overnight at 4℃, then treated with HRP-conjugated goat anti-rabbit IgG or goat anti-mouse IgG at room temperature for an hour. Chemiluminescent detection of HRP reaction was performed by using Immobilon Forte Western HRP substrate (#WBLUF0500) and filmed by ChemiDoc™ MP Imaging System (Bio-Rad Serial No.734BR3010).

### Mass Spectrometry

BAT was homogenized in RIPA buffer; the supernatant was collected after centrifugation at 12000g for 10 minutes at 4℃. Anti-UCP1 antibody (Cat. #14670; 1:200) was added to the supernatant and incubated overnight. Anti-UCP1 antibody was purified by A/G beads, washed at least three times, and mixed with Laemmli Sample buffer. The total sample was run with gel electrophoresis, and the SDS page was stained with Instant Blue. The SDS-PAGE was separated into five samples, subjected to in-gel trypsin digestion (Protocol: https://msf.ucsf.edu/protocols.html), and then analyzed by mass spectrometry. Briefly, wash the gel slice in 50 mM NH_4_HCO_3_. Next, wash the gel slice in a 50:50 mixture of 50 mM NH_4_HCO_3_ and CH_3_CN. Follow this with a wash in 100% CH_3_CN, then SpeedVac the gel slice until dry. Add 10 mM TCEP in 50 mM NH_4_HCO_3_ and incubate at 55°C for one hour. Subsequently, add 50 mM iodoacetamide in 50 mM NH_4_HCO_3_ and incubate in the dark for 30 minutes. Wash the gel slice again in 50 mM NH_4_HCO_3_, then in 100% CH_3_CN, and SpeedVac for 5 minutes. Add 20 ng/µL trypsin to completely cover the gel slices and incubate at 37°C overnight. Add Milli-Q water and transfer the supernatant to a new tube. Rinse the original tube with 50% CH_3_CN with Formic Acid, transferring the supernatant to the new tube; repeat this step three times. Finally, wash the tube with 100% CH_3_CN and transfer the supernatant to the new tube. SpeedVac the new tube until dry. The resulting pellet is now ready for mass spectrometry analysis. The procedures and data analysis were done in the Mass Core facilities of the Genomic Research Center, Academia Sinica. 129 proteins were identified by mass spectrometry (Table 1).

**Table 1.**
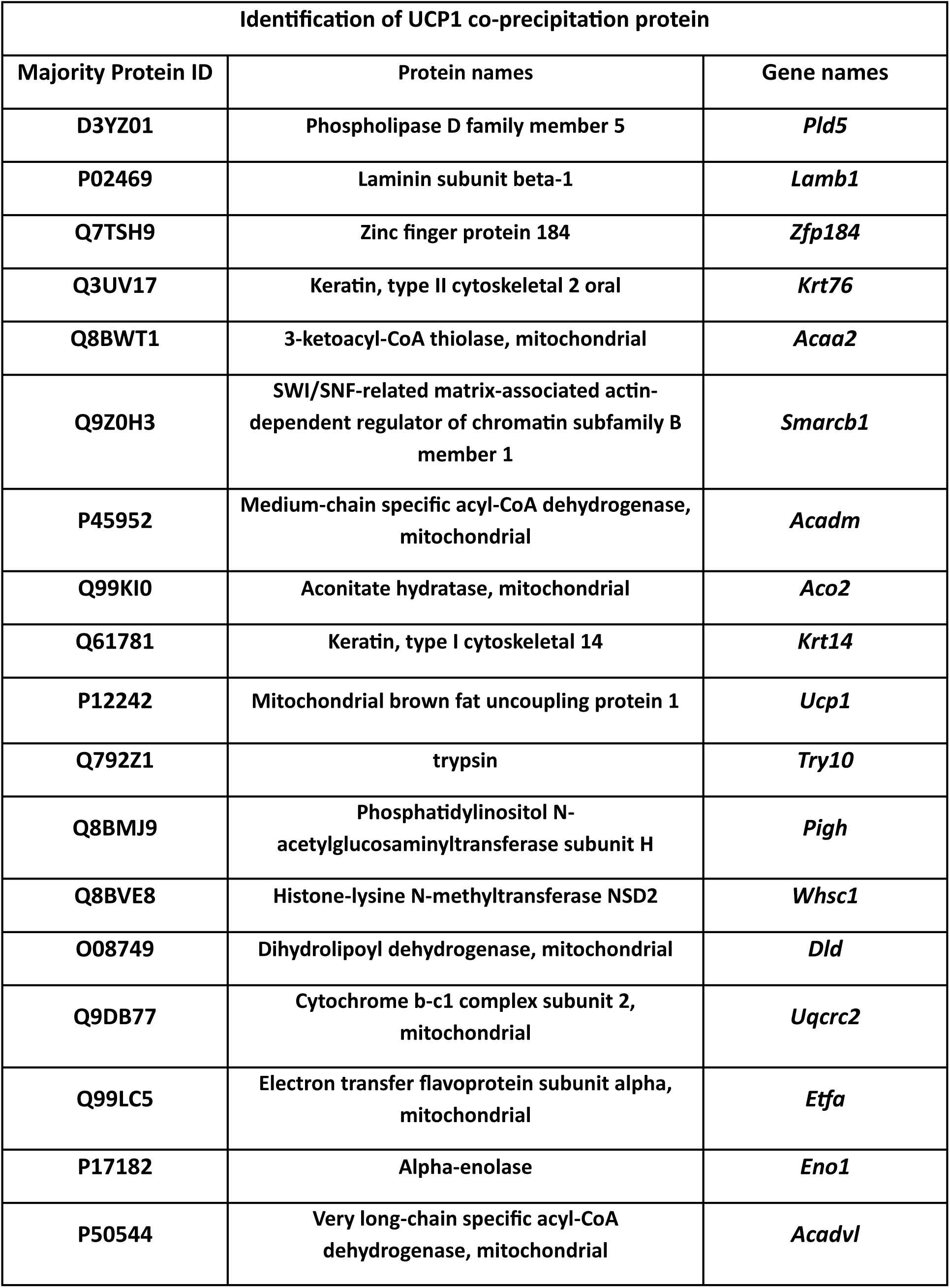
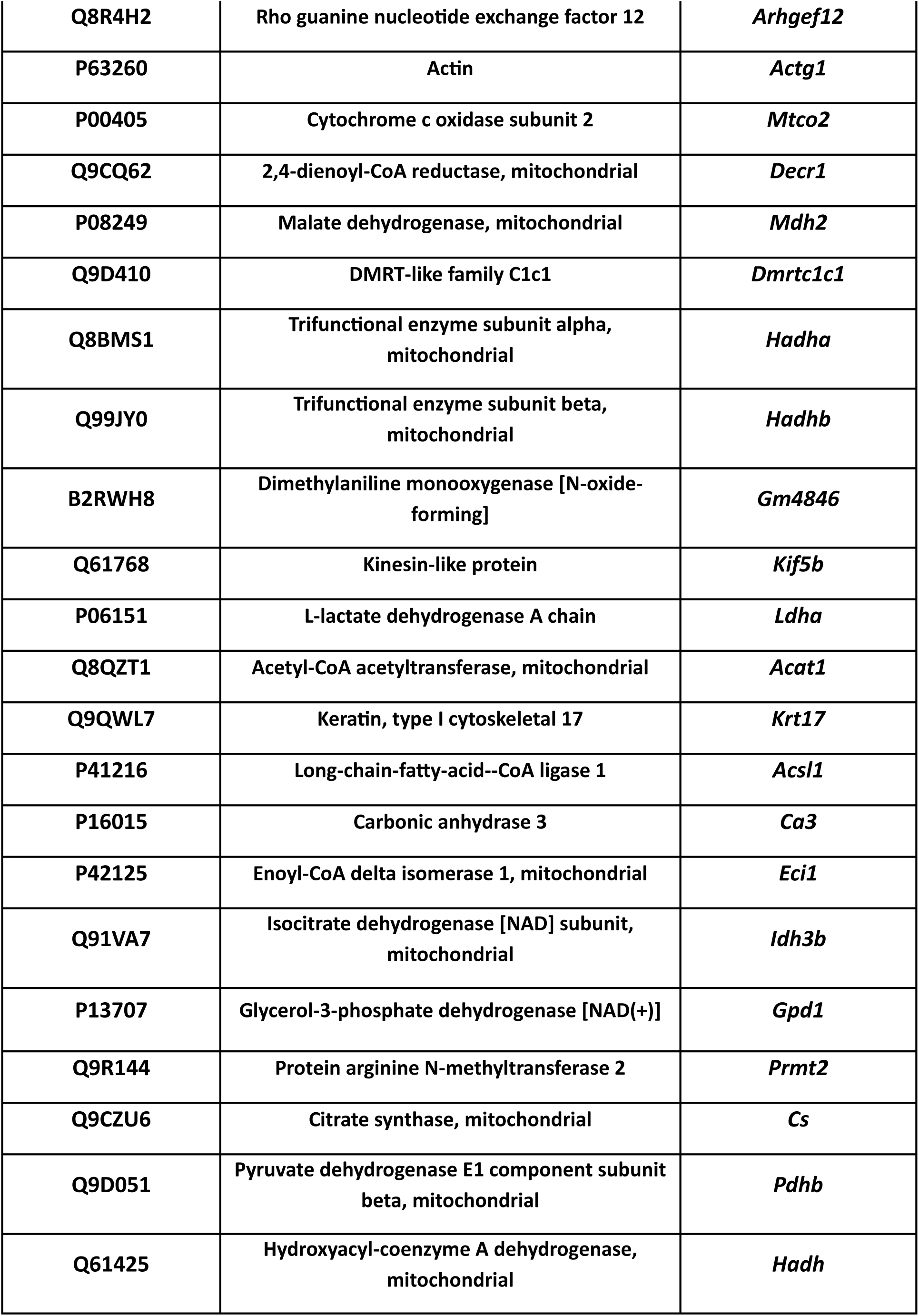
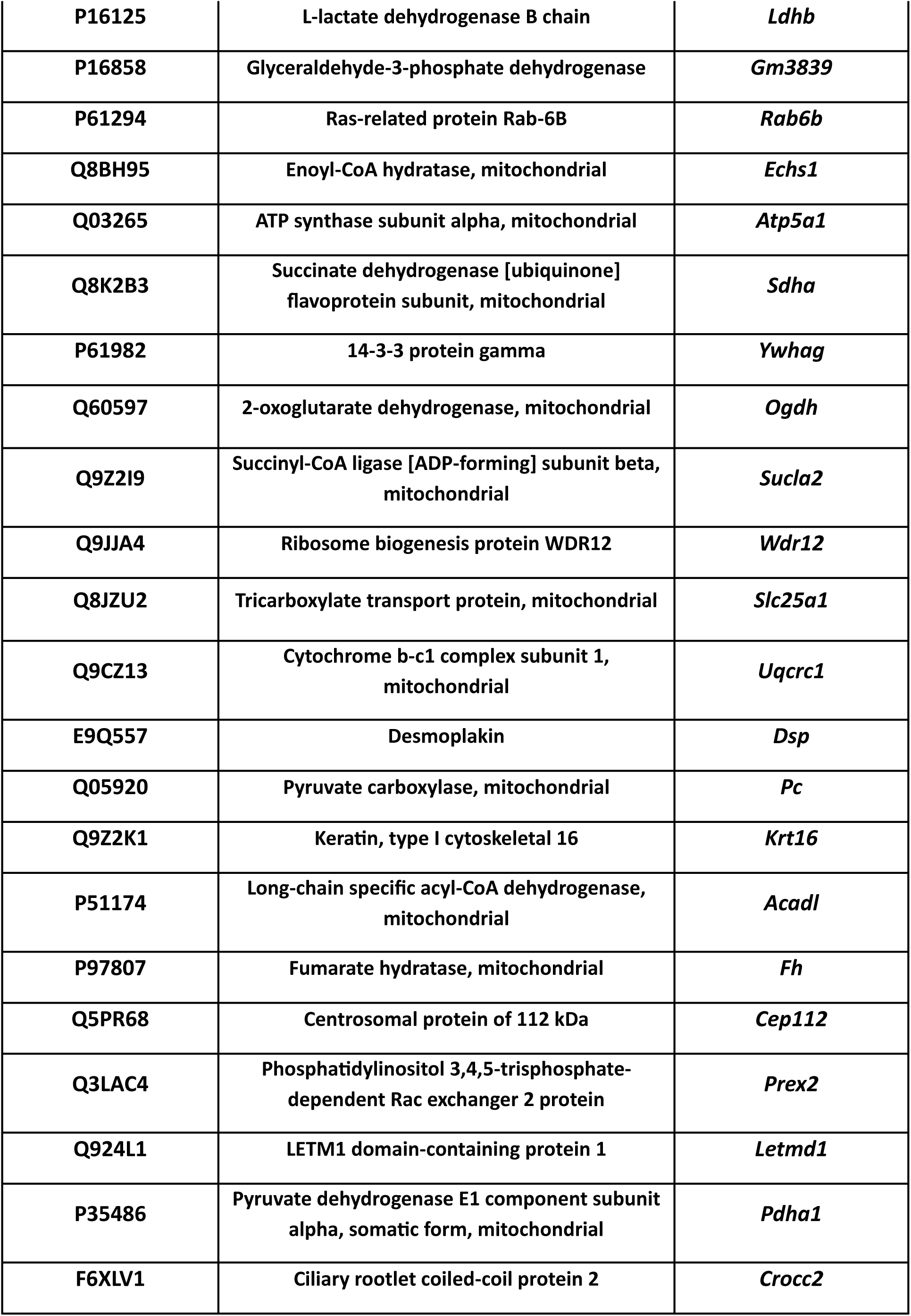
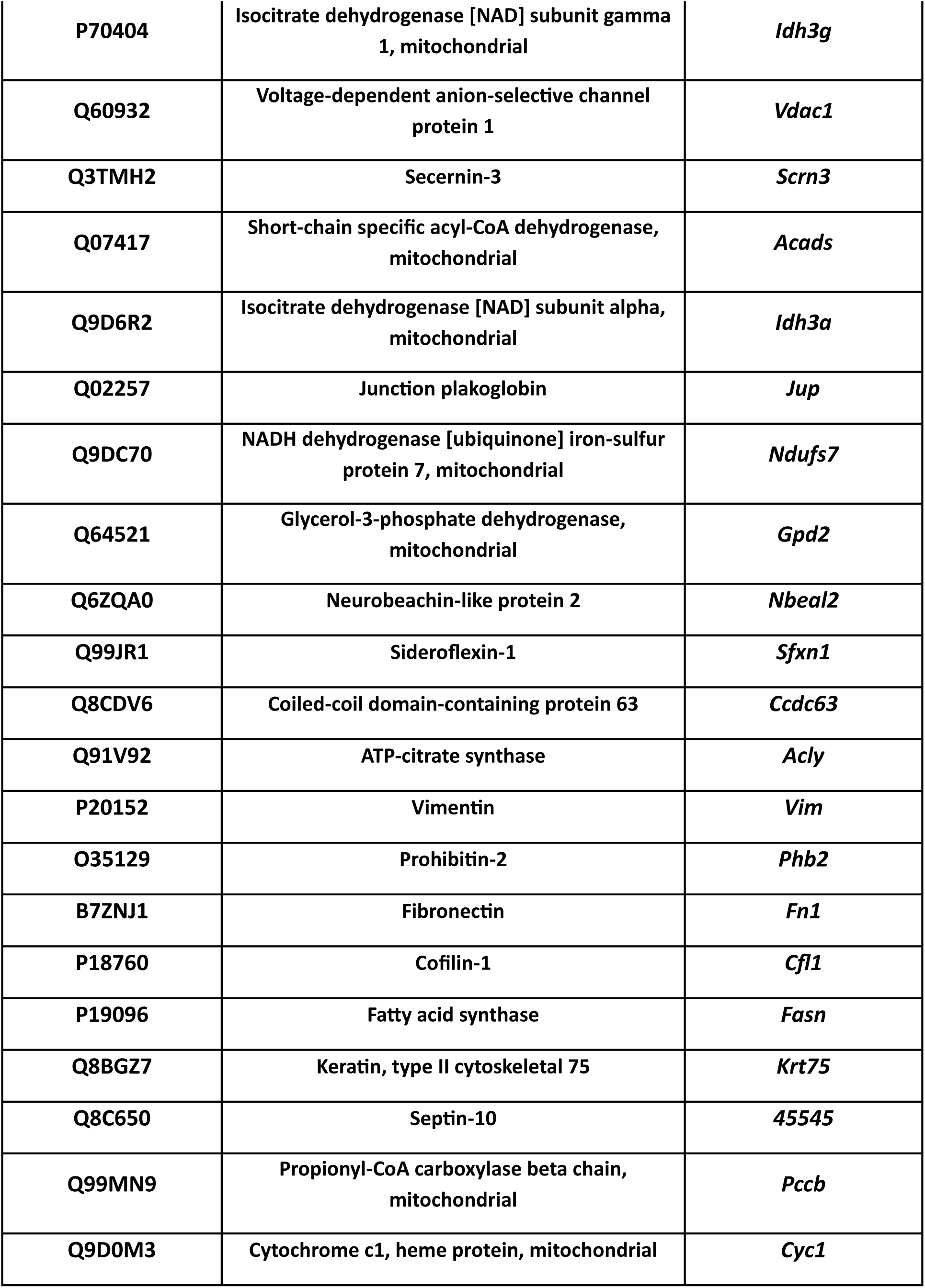
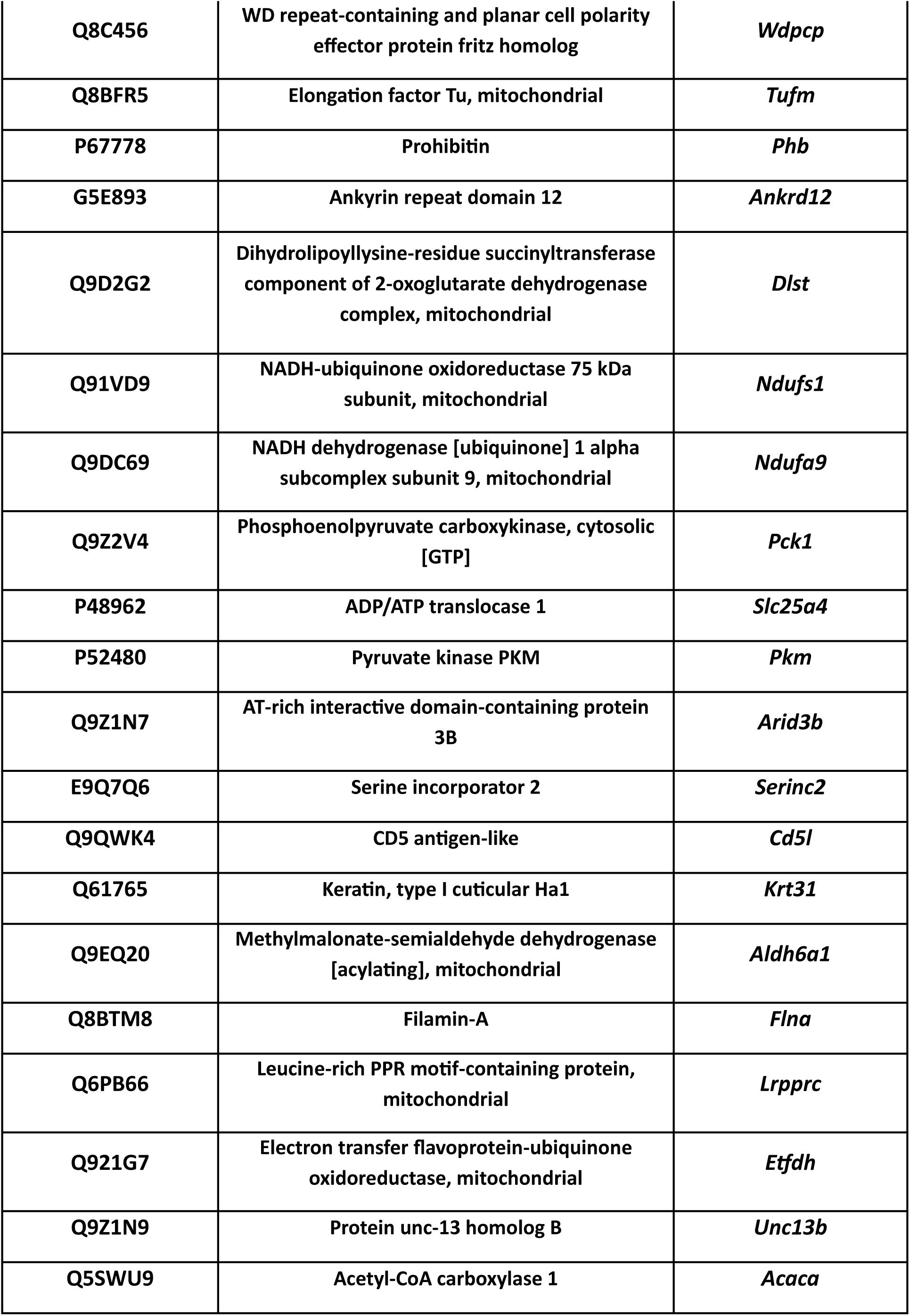
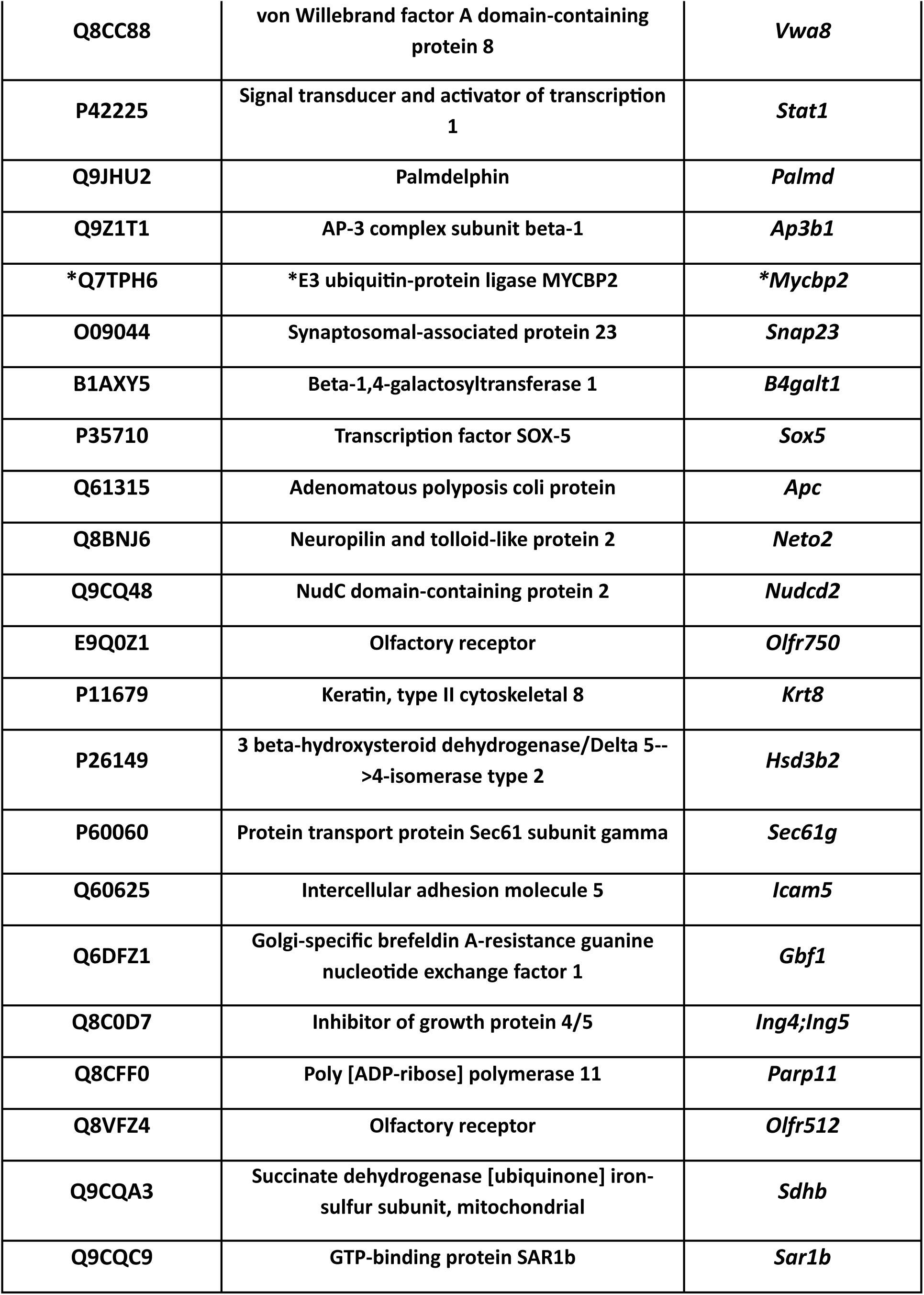
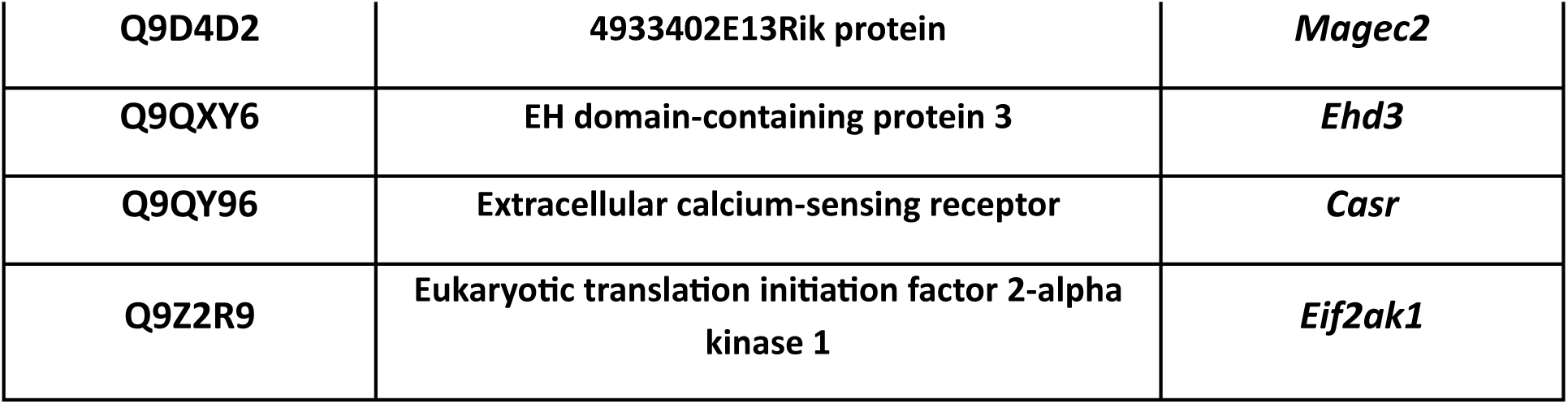
Identification of the UCP1 co-precipitation protein.

### Hudson Fixation Index (F_ST_)

To estimate the population differentiation of SNP rs1057001, we perform the Fixation Index (F_ST_) based on Hudson, R R et al.(Hudson *et al*, 1992). The figure was drawn by Python. The formulas are as follows:

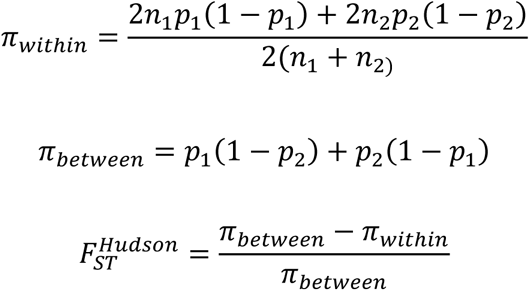

### Dual luciferase activity assays

WT1 cells were transfected with 0.5μg Dual-Glo luciferase reporter plasmid by using Turbo transfection reagent (#R0531). After 48 hours, cells were washed once with PBS and harvested with 100µl of 1×passive lysis buffer (Promega). Firefly and Renilla luciferase activities in the cell lysates were measured with the Dual-Luciferase reporter assay system (#E2940) according to the manufacturer’s protocol. Luminometry readings were obtained using a luminometer, Molecular Devices SpectraMax iD5.

### Organization, Quantification, and Statistical Analysis

Biological replicates were conducted for all WT1 cell-related experiments. Quantification of Western blot analysis was performed using GEL-Pro software. The statistical analysis was performed using Excel and Prism 10. Symbols *, **, ***, and **** indicate significance levels of *P* < 0.05, *P* < 0.01, *P* < 0.001, and *P* < 0.0001, respectively. The proposed model was drawn using Biorender or Sketchbook on an iPad. Images were organized using Adobe Photoshop and InDesign.

## The Paper Explained

### PROBLEM

Obesity and related metabolic disorders remain major global health concerns, mainly driven by an imbalance between energy intake and expenditure. Adaptive thermogenesis in brown adipose tissue (BAT), regulated by uncoupling protein 1 (UCP1), plays a central role in energy dissipation.

However, the molecular mechanisms controlling UCP1 stability and activity are incompletely understood. Understanding how UCP1 is regulated could reveal new therapeutic targets to enhance thermogenesis and combat obesity.

### RESULT

We identified a human TRIB2 variant whose frequency correlates with latitude and environmental temperature, suggesting an adaptive role in thermogenesis. Functional studies revealed that loss of Trib2 in mice protects against diet-induced obesity, alleviates hepatic steatosis, and improves glucose tolerance and insulin sensitivity. Mechanistically, TRIB2 acts as a scaffold that recruits the E3 ligase MYCBP2 to promote ubiquitination and proteasomal degradation of UCP1. Trib2 deficiency stabilizes UCP1 protein, leading to elevated thermogenic capacity and enhanced heat production in BAT.

### IMPACT

Our findings uncover a previously unrecognized post-translational mechanism that regulates UCP1 stability and thermogenesis. By linking TRIB2 function to adaptive energy expenditure and metabolic health, this study identifies TRIB2 as a potential therapeutic target for obesity and metabolic diseases. Modulating TRIB2 activity may provide a strategy to increase thermogenic efficiency and counteract metabolic imbalance.

## Data and materials availability

All data needed to evaluate the conclusions in the paper are presented in the paper and/or the Appendix. The data can be provided by the owner of the data, pending scientific review and a completed material transfer agreement. Requests for the data should be submitted to:

**Expanded View 1. Knockout of *Trib2* in mice does not affect growth**.

(A&B) The schematic diagram of generating *Trib2* KO mice and PCR genotyping of *Trib2* mice.

(C) The head and femur length were not different between *Trib2* WT and KO mice, suggesting that the knockout of *Trib2* does not affect the growth of mice.

Data C was analyzed by two-way ANOVA, followed by Tukey’s post hoc multiple comparisons test. All data represent the mean ± SEM. \**p* < 0.05, \*\**p* < 0.01; \*\*\**p* < 0.001; \*\*\*\**p* < 0.0001.

**Expanded View 2. Knockout of *Trib2* in mice attenuates hepatic steatosis and improves blood glucose homeostasis**.

(A) *Trib2* KO mice alleviate the hepatocellular steatosis compared to WT mice when fed a HFHSD.

(B) The hepatic TG was significantly reduced in *Trib2* KO mice.

(C&D) The *Trib2* KO mice fed a HFHSD significantly decrease blood glucose in the i.p.GTT and i.p.ITT test, suggesting that *Trib2* KO mice have good glucose clearance and insulin sensitivity.

(E) Serum levels of triglyceride and adiponectin were reduced in the *Trib2* KO mice fed a HFHSD.

(F) F4/80 IHC staining observed a Crown-like structure in WT mice fed with HFHSD.

Data B was analyzed by one-way ANOVA, followed by Tukey’s post hoc multiple comparisons test. Data C and D, blood glucose data, were analyzed using two-way ANOVA, followed by Šídák’s post hoc test for multiple comparisons; The AUC and inverse AUC data were analyzed using one-way ANOVA, followed by Tukey’s post hoc test for multiple comparisons. Data E was analyzed by student unpaired t-tests. All data represent the mean ± SEM. \**p* < 0.05, \*\**p* < 0.01; \*\*\**p* < 0.001; \*\*\*\**p* < 0.0001.

**Expanded View 3. The food consumption and the indirect calorimetry of *Trib2* WT and KO mice on a HFHSD**.

(A) The food consumption was not different between *Trib2* WT and KO mice.

(B) The running wheel physical activity test was not different between *Trib2* WT and KO mice. Data A and B were analyzed by student unpaired t-tests. All data represent the mean ± SEM. \**p* < 0.05, \*\**p* < 0.01; \*\*\**p* < 0.001; \*\*\*\**p* < 0.0001.

**Expanded View 4. Knockout of *Trib2* in mice does not promote white fat browning**.

(A) Immunohistochemistry staining of UCP1 on perigonadal fat.

(B) UCP1 was present in brown adipose tissue (BAT) and did not express in inguinal adipose tissue (iWAT) and gonadal adipose tissue (gWAT).

**Expanded View 5. TRIB2 knockdown enhances UCP1 expression and mitochondrial function in WT1 Cells**.

(A) Knockdown of TRIB2 increases UCP1 expression in knockdown TRIB2 Day 8 differentiation WT1 cells.

(B) The mitochondrial function was improved in knockdown WT1 cells.

Data was analyzed by student unpaired t-tests. All data represent the mean ± SD. \**p* < 0.05, \*\**p* < 0.01; \*\*\**p* < 0.001; \*\*\*\**p* < 0.0001.

